# Quantitative Cytogenetics Reveals Molecular Stoichiometry and Longitudinal Organization of Meiotic Chromosome Axes and Loops

**DOI:** 10.1101/724997

**Authors:** Alexander Woglar, Kei Yamaya, Baptiste Roelens, Alistair Boettiger, Simone Köhler, Anne M Villeneuve

## Abstract

During meiosis, chromosomes adopt a specialized organization involving assembly of a cohesin-based axis along their lengths, with DNA loops emanating from this axis. We applied novel, quantitative and widely applicable cytogenetic strategies to elucidate the molecular bases of this organization using *C. elegans*. Analyses of WT chromosomes and *de novo* circular mini-chromosomes revealed that meiosis-specific HORMA-domain proteins assemble into cohorts in defined numbers and co-organize the axis together with two functionally-distinct cohesin complexes (REC-8 and COH-3/4) in defined stoichiometry. We further found that REC-8 cohesins, which load during S phase and mediate sister chromatid cohesion, usually occur as individual complexes, supporting a model wherein sister cohesion is mediated locally by a single cohesin ring. REC-8 complexes are interspersed in an alternating pattern with cohorts of axis-organizing COH-3/4 complexes (averaging three per cohort), which are insufficient to confer cohesion but can bind to individual chromatids, suggesting a mechanism to enable formation of asymmetric sister chromatid loops. Indeed, immuno-FISH assays demonstrate frequent asymmetry in genomic content between the loops formed on sister chromatids. We discuss how features of chromosome axis/loop architecture inferred from our data can help to explain enigmatic, yet essential, aspects of the meiotic program.

## INTRODUCTION

At the onset of meiotic prophase in organisms ranging from yeast to humans, newly replicated chromosomes adopt a highly specialized organization that enables diploid cells to produce haploid gametes. This reorganization involves the assembly of a discrete axial structure along the lengths of each conjoined sister chromatid pair, with the majority of DNA organized into loops emanating from this axis. These meiotic chromosome axes and/or their constituent components contribute to virtually all major aspects of meiotic prophase, including chromosome movement, homologous chromosome pairing, formation of DNA double strand breaks (DSBs), assembly of Synaptonemal Complex (SC) between aligned homologs, and repair of DSBs by interhomolog recombination to yield crossovers that will ensure homolog segregation. Thus, understanding the structure and function of meiotic chromosome axes is crucial for elucidating the mechanisms responsible for faithful inheritance of chromosomes during sexual reproduction.

A combination of genetic, cytological and biochemical approaches has identified the major components of meiotic chromosome axes in multiple different experimental systems. The foundation of the chromosome axis is built up by meiosis-specific cohesin complexes that are composed of a mixture of canonical subunits and meiosis-specific subunits. Most notably, meiosis-specific kleisin subunits have been identified in multiple organisms (Ishiguro, 2019), including Rec8p in budding yeast (Klein et al., 1999)), RAD21L and REC8 in mammals (Eijpe et al., 2003; Parisi et al., 1999), and COH-3, COH-4 and REC-8 in *C. elegans* (Pasierbek et al., 2001; Severson et al., 2009). At the onset of meiotic prophase, these cohesin complexes mediate by unknown mechanisms the recruitment of other meiosis-specific axis components. Some of these, including SYCP2/SYCP3 in mouse (Eijpe et al., 2000; Schalk et al., 1998) and Red1p in budding yeast (Smith and Roeder, 1997) have been designated as “axis core” proteins, as they play a role in recruitment of meiosis-specific HORMA-domain containing proteins (*e*.*g*. Hop1p in yeast (Hollingsworth and Byers, 1989) or HORMAD1/2 in mammals (Wojtasz et al., 2009); referred to as “HORMADs” from here on). The HORMADs share a structural organization in common with spindle-assembly checkpoint protein Mad2, which has a capacity to form complexes with other proteins through topological entrapment of peptides known as “closure motifs” by a Mad2 domain known as a “safety belt” (Rosenberg and Corbett, 2015). Recent work has demonstrated that yeast Red1p and mammalian SYCP-2/SYCP-3 each contain a single closure motif that mediates interactions with their respective HORMAD partners; further, Red1p homo-tetramers and SYCP2/SYCP3 hetero-tetramers are capable of forming oligomeric filaments *in vitro* (West et al., 2019). In *C. elegans*, similar axis core components have not been identified, but four different HORMAD paralogs are present and constitute a hierarchical complex that builds up the meiotic chromosome axis (HTP-3, HTP-1/2 and HIM-3; (Couteau and Zetka, 2005; Goodyer et al., 2008; Martinez-Perez and Villeneuve, 2005; Martinez-Perez et al., 2008; Zetka et al., 1999). Further, biochemical and *in vivo* experiments have demonstrated that HTP-3 recruits HTP-1/2 and HIM-3 to the axis by interacting with their HORMA domains via closure motifs located in its C-terminal tail (Kim et al., 2014). Another recent study provided information regarding the relative cross-sectional positioning of cohesins and HORMADs within the context of the mature SCs of *C. elegans*, showing that HORMADs are located closer to the central region of the SC, whereas cohesin complexes are located closer to the bulk of the chromatin (Kohler et al., 2017).

While there has been substantial progress in identifying components of the meiotic chromosome axis and interactions among many of these components, much remains to be learned regarding how these proteins and interactions become organized along the length of chromosomes into a functional composite structure that mediates and coordinates key meiotic events. Ensemble/population-based measurements have identified preferential sites of association of multiple meiotic axis proteins in budding and fission yeast (Miyoshi et al., 2012; Panizza et al., 2011), but these analyses do not address how many (and which) sites are occupied at the same time on individual meiotic chromosomes and in a given individual meiocyte. Further, a recent Hi-C-based study of mouse spermatocytes suggests a lack of reproducible loop positions during mouse meiosis (Patel et al., 2019). It is also not known (either in meiotic cells or mitotically-dividing cells) how many cohesin complexes are required to locally provide sister chromatid cohesion (“single/simple ring” *vs*. “handcuff model”, e.g.: see (Rankin and Dawson, 2016)), or how many cohesins and other axis forming proteins load onto a given stretch of DNA and organize a pair of sister chromatids into a linear axis with emanating chromatin loops, as they appear in cytological preparations. Consequently, we currently lack understanding regarding how meiotic chromosome organization accomplishes certain essential tasks. For example: programmed meiotic DSBs must use the homologous chromosome rather than the sister chromatid as a repair template for recombinational repair, as it is necessary to form a crossover between homologs to provide the basis for a temporary link between the homologs that will ensure their correct segregation at the first meiotic division. An “*inter-sister block*” or “inter-*homolog bias*” favoring use of the homolog as a recombination partner is a long-known phenomenon in the meiotic recombination program and depends on chromosome axis proteins (for review see: (Zickler and Kleckner, 2015)), but how this bias is achieved is not understood on a mechanistic level. It is clear that our knowledge regarding how meiotic chromosome structures confer characteristic properties of the meiotic program would benefit from a fuller understanding of how the axis itself is organized.

Here, we report an analysis of the molecular stoichiometry and longitudinal architecture of the meiotic chromosome axis, using the well-established meiotic model organism *C. elegans*. Our work builds on and exploits a well-recognized feature of meiotic chromosomes from various model (and non-model) systems, namely that when meiotic nuclei are strongly spread out in 2D on a glass slide (“fully spread” from here on), the continuous axis configuration (observed in in situ preparations and under mild/partial spreading conditions) is “spaced out”, and cohesins, axis core proteins and/or HORMADs can appear as linear arrays of foci visible by electron microscopy or immuno-fluorescence (*e*.*g*. see: (Anderson et al., 1988; Ishiguro et al., 2011; Kim et al., 2010; Smith and Roeder, 1997) and FIG1A). By combining this spreading approach with novel quantitative strategies and/or structured illumination microscopy (SIM), we reveal previously unrecognized features of meiotic chromosome organization. We demonstrate that different HORMAD proteins assemble into cohorts of defined numbers and co-organize the chromosome axis together with two structurally and functionally distinct cohesin complexes in defined stoichiometry. We show that half of the REC-8-containing cohesin complexes loaded during S phase are abruptly lost concurrent with loading of COH-3/4 cohesin complexes and axis assembly. In the resulting axis, individual cohesion-mediating REC-8 complexes occur on average every 130-200 kbp, interspersed with cohorts containing an average of three axis-organizing COH-3/4 complexes and three HORMAD complexes. Finally, we provide evidence that the axis architecture deduced from our analysis is associated with an inherent asymmetry in size, number and genomic composition between the loops formed on sister chromatids. Together, our analyses provide a quantitative framework that will inform and constrain future experiments and models regarding meiotic chromosome axis organization and function. Moreover, the quantitative cytogenetic strategies applied here should be broadly applicable for investigating molecular stoichiometry in the context of other cellular structures and processes.

**Figure 1.**
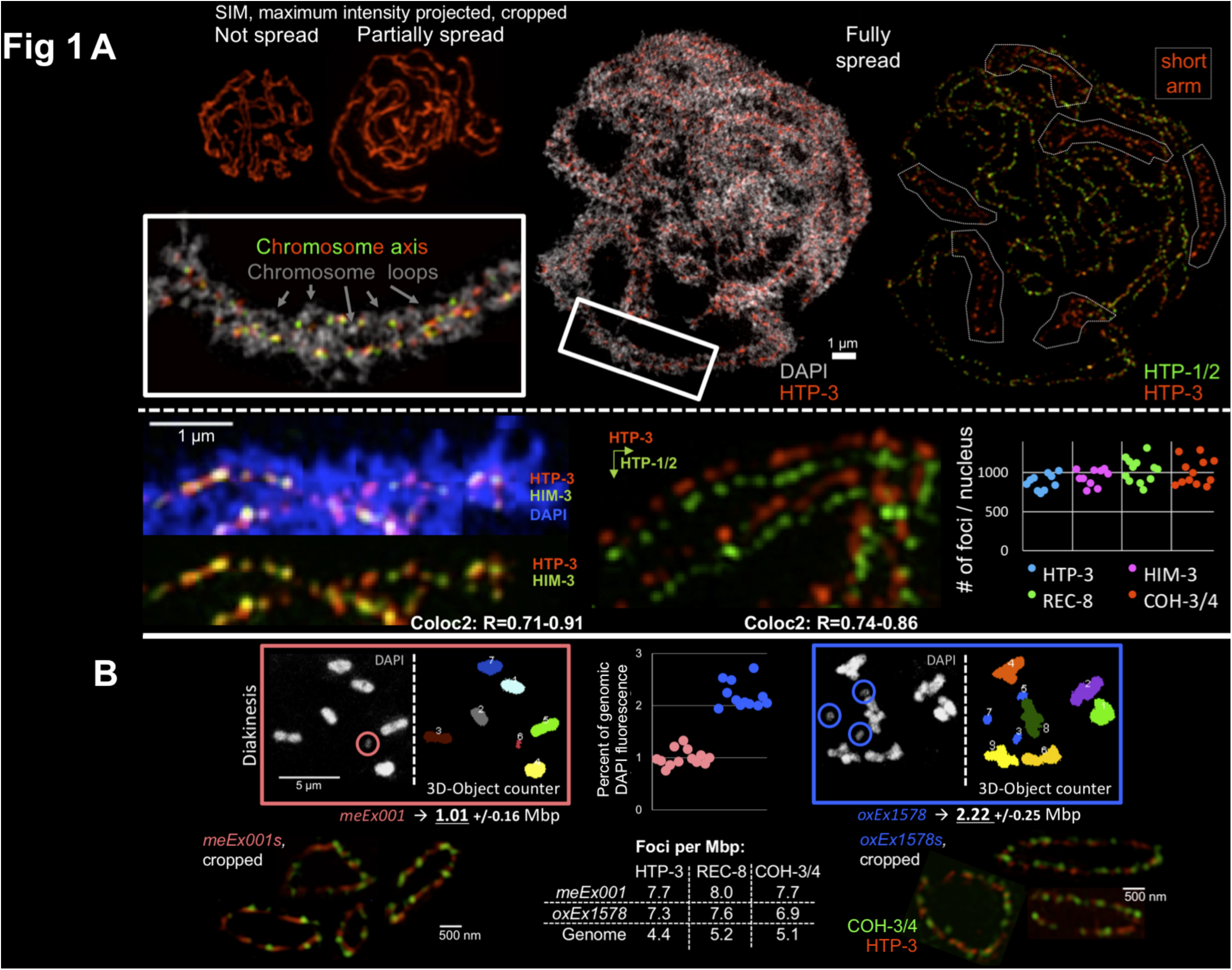
Components of the chromosome axis can be resolved as individual foci. A) TOP: Comparison of Structured Illumination Microscopy (SIM) images of *in situ*, partially spread, and fully spread individual meiotic prophase nuclei, displayed at the same magnification. In the image at the right, dashed lines outline the domains of each of the six bivalents where HTP-1/2 proteins have become depleted from the chromosome axis, indicating that this nucleus is at the late pachytene stage. BOTTOM LEFT, CENTER: Images of cropped segments of synapsed chromosome pairs, illustrating both the pearls-on-a-string appearance of chromosome axis foci and colocalization of HTP-3 and HIM-3 (overlaid) or HTP-1/2 (images offset in both x and y); range of R values from ImageJ Coloc2 analysis for 8-10 individual fully spread nuclei. BOTTOM, RIGHT: Quantification of numbers of HTP-3, HIM-3 REC-8 and COH-3/4 foci in individual fully spread nuclei; each data point represents a nucleus. B) TOP: Method used to estimate the total DNA content of two distinct ExChrs, *meEx001* and *oxEx1578*. Sample deconvolved wide-field images of DAPI-stained chromosome complements from individual oocyte nuclei (with the ExChrs circled) are presented together with masks generated using the “3D Object Counter” ImageJ plugin; in the accompanying graph, each data point represents the total DAPI fluorescence for a single ExChr (2C), normalized to ½ of the total DAPI fluorescence measured for all 6 bivalents (4C) in the same nucleus. BOTTOM, LEFT and RIGHT: Individually cropped examples of ExChr *meEx001* (left) and *oxEx1578* (right) from fully spread nuclei stained for COH-3/4 or REC-8 together with HTP-3. BOTTOM, CENTER: Table representing the average densities per Mbp of HTP-3, REC-8 and COH-3/4 foci derived from our analyses counting foci for the whole genome and for the ExChrs (see FIG1S2B and FIG1A, bottom).

## RESULTS and DISCUSSION

### The meiotic chromosome axis is a linear array of discrete cohorts of cohesins and HORMADs

In contrast to the more continuous appearance of HORMAD signals observed in *in situ* preparations of *C. elegans* gonads and under partial spreading conditions (which maintain the spatial-temporal organization of the gonad largely intact), we find that in fully spread *C. elegans* meiotic prophase nuclei, the chromosome axis is resolved as individual HORMAD foci, arranged like pearls on a string (FIG1A, FIG1S1A), as observed in other organisms. The three types of HORMAD proteins that constitute the *C. elegans* meiotic chromosome axis (HTP-3, HTP-1/2, and HIM-3; (Goodyer et al., 2008; Martinez-Perez et al., 2008; Zetka et al., 1999)) colocalize together in most of these foci in early prophase nuclei when examined by Structured Illumination Microscopy (SIM, FIG1A), consistent with the demonstrated direct physical interactions among these components (Goodyer et al., 2008; Kim et al., 2014). Further, previously reported findings concerning distinct behaviors of different HORMAD proteins could be recapitulated in these fully spread preparations. For example, in late prophase, crossover designation triggers the reorganization of the *C. elegans* meiotic bivalent into two distinct domains: a long arm, where all the HORMADs are present, and a short arm, where only HTP-3 and HIM-3 are present (Martinez-Perez et al., 2008); this reorganization is clearly visible in the fully spread preparations (FIG1A). We count approximately 1000 HORMAD-containing foci (HIM-3 and HTP-3) in fully spread pachytene nuclei (FIG1A, bottom right).

*C. elegans* HORMAD protein HTP-3, which is required to load all the other HORMADs (Goodyer et al., 2008), appears to play a role in axis organization analogous to the axis core proteins (Red1 and SYCP2 and 3) identified in other organisms. In the absence of HTP-3, meiotic cohesin complexes still bind chromatin and hold sister chromatids together until anaphase (Severson et al., 2009), but they are not arranged as a continuous axis or in a pearls-on-a-string-like configuration during prophase (FIG1S1). Thus, absence of HTP-3 phenocopies budding yeast *red1* mutants (in which axes don’t form, but cohesin binding is unaffected), rather than *hop1* mutants (in which Red1p and Rec8p containing axes form) (Klein et al., 1999; Woltering et al., 2000). Therefore, we speculate that HTP-3, which contains six closure-motifs that mediate associations with HIM-3 and HTP-1/2 (Kim et al., 2014), functions as the axis core component in *C. elegans*.

During metazoan meiosis, chromosome axis organization and meiotic sister chromatid cohesion are mediated by at least two different cohesin complexes that contain distinct kleisin subunits and/or load at distinct times. Sister chromatid cohesion is mediated by complexes containing REC-8 in worms, REC8 in mammals and C(2)M in flies, whereas axis formation requires COH-3 and COH-4 in worms, REC8 and RAD21L in mammals and the SOLO-SUNN-containing cohesin complex in flies (Crawley et al., 2016; Gyuricza et al., 2016; Pasierbek et al., 2001; Severson et al., 2009; Xu et al., 2005). Because COH-3 and COH-4 are products of a recent gene duplication, are functionally redundant and are recognized by the same antibody, they are referred to as COH-3/4 from here on (Severson et al., 2009; Severson and Meyer, 2014). We count similar numbers of foci for both REC-8 cohesin complexes and COH-3/4 cohesin complexes as we count HORMAD protein foci (about 1000 foci per nucleus, FIG1A). If we consider the numbers of HORMAD foci, REC-8 foci or COH-3/4 foci to be distributed along the diploid genome (2 x 100 Mbp), we can estimate an average of one focus (of each type) every ∼200 kbp.

Because overlap between chromosome segments in spreads of whole nuclei might result in an underestimate of foci numbers, we sought to validate this estimate by measuring numbers of foci on well-separated DNA segments of defined size. Our approach was to evaluate numbers of cohesin and HORMAD foci on extra-chromosomal arrays (ExChrs), which are mini-chromosomes that form via fusion of linear or circular DNA of any origin upon injection into the *C. elegans* gonad. ExChrs are visible as additional DAPI bodies in oocytes, which allows an approximation of their size (FIG1B). Depending on the mixture of injected DNAs, ExChrs can be highly repetitive or complex in DNA sequence. Repetitive ExChrs acquire heterochromatic marks and are silenced in the germ line, whereas complex ExChrs can be transcriptionally active in germ cell nuclei (Kelly et al., 1997). ExChrs can be transmitted through mitosis and can be inherited across generations, albeit in non-Mendelian fashion (Mello et al., 1991; Stinchcomb et al., 1985).

Here we analyzed ExChr *meEx001*, a complex array that has 95% *E. coli* genomic DNA, and *oxEx1578*, a repetitive array that has 0% *E. coli* genomic DNA. Based on 3D DAPI intensity measurements in diakinesis-stage oocytes, we estimate that *meEx001* contains about 1 Mbp of DNA and *oxEx1578* contains about 2.2 Mbp, corresponding to approximately 1% and 2% of the size of the *C. elegans* haploid genome, respectively (FIG1B). Interestingly, we found that ExChrs are ring chromosomes (FIG1S2A) as suggested by S.K. Kim in 1992 (http://wbg.wormbook.org/wli/wbg12.2p22). Both *meEx001* and *oxEx1578* form chromosome axes during meiotic prophase, but do not engage in synapsis when more than one ExChr is present in the nucleus (FIG1S2B).

In fully spread nuclei, we find that ExChrs are decorated by HTP-3, COH-3/4 and REC-8 foci in a pattern similar to that observed on normal chromosomes (FIG1B). Numbers of HORMAD foci and cohesin foci on a given ExChr are strongly correlated (FIG1S2B). Further, the average numbers of foci observed for these two ExChrs were proportional to the amount of DNA present, as we detected twice the number of foci on the ExChr that contains twice the amount of DNA (*meEx001*: HTP-3: 7.8(+/-2.4), REC-8: 8.1(+/-2.5) and COH-3/4: 7.8(+/-2.6) foci; *oxEx1578*: HTP-3: 16.1(+/-2.9), REC-8: 16.8(+/-2.8) and COH-3/4: 15.3(+/-3.1) foci, FIG1C and FIG1S2B).

Our observation that *meEx001* and *oxEx1578* exhibit the same average numbers of cohesin and HORMAD foci (∼1 focus of each type per 130 kbp of DNA) is interesting given that these two ExChrs share almost no sequences in common with each other or with the *C. elegans* genome (except for a promoter fragment and plasmid sequences in the selection markers that are present in vastly different amounts in the two ExChrs). Thus, we conclude that HORMAD, REC-8 and COH-3/4 foci occur in similar numbers along chromosomal DNA as a function of chromosomal size and that they can do so apparently independently of DNA sequence.

### Non-random arrangement of structurally and functionally distinct cohesin complexes along chromosome axes

Previous work provided evidence that REC-8 and COH-3/4 cohesin complexes play functionally distinct roles during *C. elegans* meiosis (Severson et al., 2009; Severson and Meyer, 2014): REC-8 has been demonstrated to mediate sister chromatid cohesion, whereas COH-3/4 plays a key role in axis organization (FIG2S1A). Further, these functionally distinct complexes differ in their timing of association with chromatin: REC-8 cohesins (like murine REC8) load during pre-meiotic DNA replication, whereas COH-3/4 cohesins (like murine RAD21L) load after completion of S phase, at the beginning of prophase ((for review: (Ishiguro, 2019)) and are insufficient to maintain connections between sister chromatids in the absence of recombination (Crawley et al., 2016). Our data support and extend these findings substantially in several ways.

First, measurement of fluorescence levels in immuno-stained gonads demonstrated that half of the REC-8 cohesin complexes loaded during S phase are removed from chromatin concurrently with loading of COH-3/4 cohesin complexes and coalescence of meiotic chromosome axes (FIG2A). This suggests that a fundamental shift in the relationships between sister chromatids may occur upon entry into meiotic prophase. The sharp drop of REC-8 molecules upon meiotic prophase entry does not depend on the presence of COH-3/4, as it occurs even in a *coh-4 coh-3* mutant (FIG2S1B). Cohesin release factor WAPL-1 may play a role in REC-8 removal, as a drop in REC-8 levels upon meiotic entry is not observed in a *wapl-1* mutant (FIG2S1B); however, interpretation of this finding is complicated by the fact that COH-3/4 cohesins load prematurely at low levels during S phase in the *wapl-1* mutant (Crawley et al., 2016), so REC-8 may not be loaded at normal levels in this mutant.

**Figure 2.**
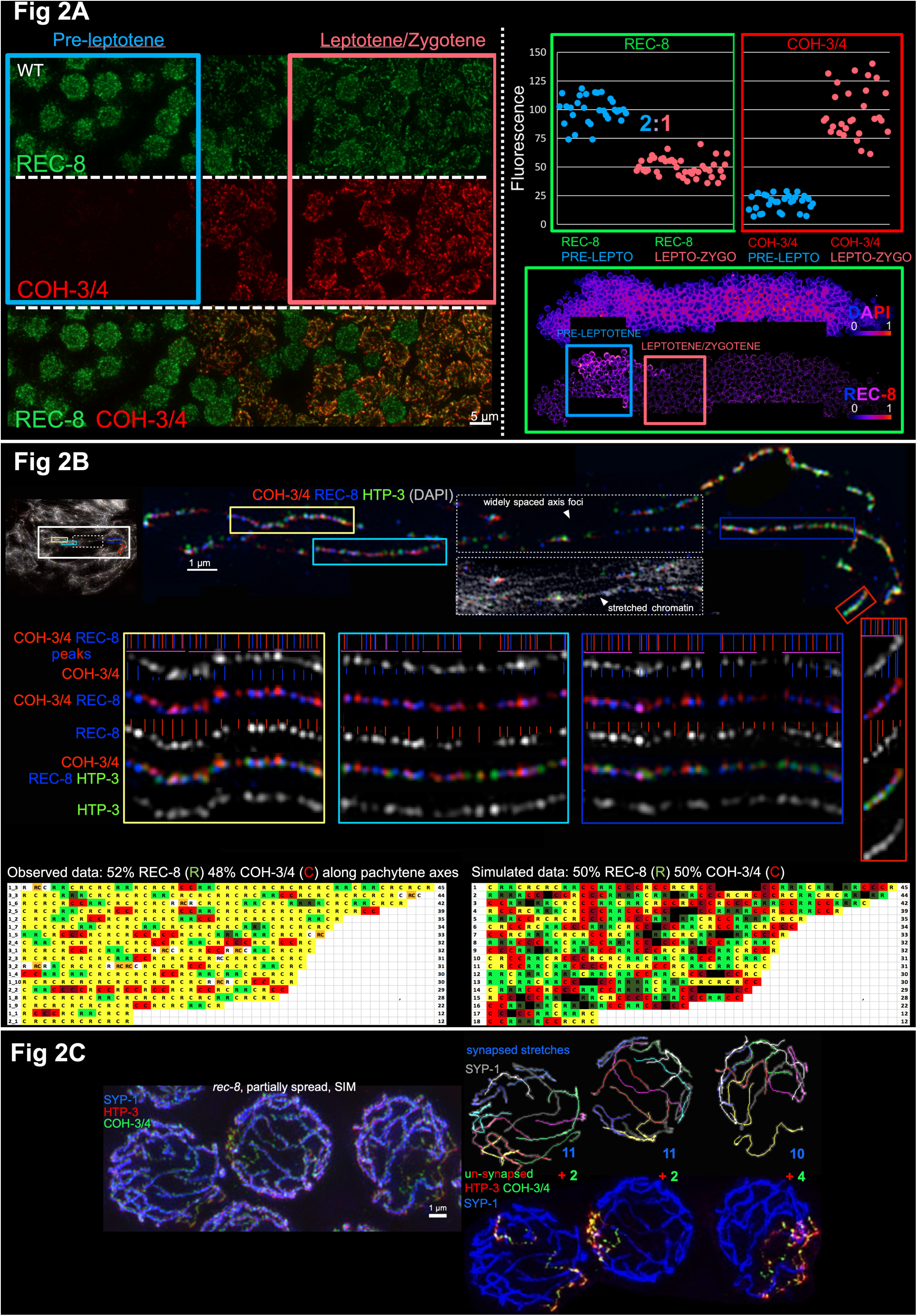
Non-random distribution of functionally and structurally distinct cohesin complexes. A) Reduction in chromosome-associated REC-8 cohesion-conferring complexes occurs concurrently with axis formation. LEFT: Deconvolved wide-field images of a partially spread gonad from a worm expressing REC-8::GFP, stained for GFP, COH-3/4 and DNA, illustrating that COH-3/4 becomes detectable on chromatin at leptotene entry and REC-8 signal intensity drops concurrently. As nuclei in this region of the gonad move approximately one row every 40 minutes (Rosu and Cohen-Fix, 2017), the fact that nuclei with medium levels of COH-3/4 or REC-8) are rarely observed suggests that this is a rapid transition. RIGHT: Quantitation of total REC-8 or COH-3/4 fluorescence in nuclei at pre-leptotene and leptotene/zygotene stages, measured using SUM projections of 3D-stiched gonads. Each data point represents a single nucleus; for each of the 4 gonads analyzed for each kleisin, fluorescence levels were normalized to 100 for the mean fluorescence in pre-leptotene nuclei for REC-8 and for the mean fluorescence in leptotene/zygotene nuclei for COH-3/4. REC-8 fluorescence drops at leptotene entry to half its pre-leptotene levels. The images below the graph illustrate the drop in REC-8 using the indicated color scale to depict fluorescence signal intensity and boxes to indicate the regions of the gonad where fluorescence was quantified. B) Non-random, alternating distribution of cohesin complexes containing different kleisin subunits along chromosome axes. TOP: SIM image of an individually cropped bivalent from a fully spread prophase nucleus, illustrating lack of colocalization of COH-3/4 and REC-8 signal peaks, even when axis segments are well preserved (colored boxes). The blue and red vertical lines in each enlarged segment represent the positions of REC-8 (blue) and COH-3/4 (red) signal peaks. Purple horizontal lines represent continuous stretches of alternating REC-8 and COH-3/4 signal peaks. BOTTOM: Each line in the in the table represents a single, Z-projected and straightened axis segment, with R indicating REC-8 foci and C indicating COH-3/4 foci. Yellow indicates that both direct neighbors of a given kleisin focus are of the other kleisin type. Red (COH-3/4) or green (REC-8) indicates that the indicated focus has one like neighbor and one different neighbor. Darkening red or green represent more than two kleisins of the same kind in a row. Orange represents two, non-resolvable REC-8 and COH-3/4 signal peaks (such as the first signal peak in top, right). Axis segments are sorted by the number of cohesin foci (indicated at the right); all analyzed segments are presented (n = 18 segments from 3 nuclei). REC-8 and COH-3/4 foci alternate with each other in a highly non-random fashion (*e.g*.: stretches of three or more of the same kleisin were observed much less frequently than expected for a random distribution (10 observed *vs*. 131 expected; p<0.00001, chi-square test, www.socscistatistics.com). The table on the right represents simulated data obtained using 50% C and 50% R as input for https://www.random.org/lists. C) Evidence for association of COH-3/4 cohesin complexes with individual chromatids. RIGHT TOP: SIM image of SYP-1, HTP-3 and COH-3/4 immuno-staining in three meiotic prophase nuclei from a *rec-8* mutant gonad; all three nuclei shown display a combination of SCs and unsynapsed chromosomes. MIDDLE: Tracings of individual SCs from the above nuclei; blue numbers indicate the total number of SCs in each nucleus. BOTTOM: Nuclei with SCs pseudo-colored in blue together with images of unsynapsed chromosome axes (marked with HTP-3 and COH-3/4); green numbers indicate the number of unsynapsed chromosome axes present in each individual nucleus. When unsynapsed chromosome stretches were detected in the *rec-8* mutant, they were consistently present in even numbers. Moreover, in nuclei where all SCs and unsynapsed axes could be reliably traced, we found that 2 x (the number of SCs) + (the number of unsynapsed axes) = 24, corresponding to the total number of chromatids present. Based on these numbers (and SC lengths measuring approximately 75-80% the lengths of normal SCs, see FIG2S1C), we infer that SCs form between sister chromatid pairs in the *rec-8* mutant and that unsynapsed chromosome stretches correspond to individual chromatids.

Second, analysis of SIM images of fully spread chromosome axes revealed that not only do COH-3/4 and REC-8 signal peaks usually not coincide, but COH-3/4 and REC-8 signal peaks usually occur in a largely alternating pattern along a given axis stretch (FIG2B). This striking alternating arrangement is significantly different from simulated random positioning along a theoretical axis, strongly suggesting a functional biological basis underlying the observed pattern (FIG2B). We note that while HORMAD protein foci were detected in numbers similar to the numbers of REC-8 and COH-3/4 foci (FIG1), we did not detect a consistent longitudinal pattern in their arrangement relative to cohesins along the axis. However, HORMAD, REC-8 and COH-3/4 foci do occur in approximately equal proportions (1:1:1) locally along the chromosome axis; further, HORMAD foci can partially overlap with REC-8 foci, COH-3/4 foci or both, albeit with their signal peaks often considerably offset (FIG2B).

Third, analysis of spread nuclei from *rec-8* mutants provides additional evidence for a functional distinction between REC-8 and COH-3/4 cohesin complexes, by demonstrating a capacity for COH-3/4 cohesin complexes to interact with individual chromatids. In *rec-8* mutants, when only COH-3/4 cohesin complexes are present on chromatin during meiotic prophase, SCs frequently form and crossover recombination apparently occurs between sister chromatids (rather than between homologs), suggesting that axes assemble along individual chromatids in this context (Crawley et al., 2016; Cahoon et al., 2019). However, not all chromosomes engage in synapsis in *rec-8* mutants (Cahoon et al., 2019), and we were able to trace the paths of both synapsed and unsynapsed axes in late prophase nuclei of *rec-8* mutants (FIG2C). This analysis confirmed the interpretation that continuous chromosome axes (containing HTP-3 and COH-3/4) do indeed form along individual chromatids in the *rec-8* mutant (FIG2C and FIG2S1C). Thus, we infer that while REC-8 cohesin complexes are required to confer cohesion between sister chromatids, COH-3/4 cohesin complexes are sufficient to organize chromosome axes, and moreover, can topologically embrace or bind to single chromatids (rather than sister chromatid pairs), at least when REC-8 cohesin is absent.

### Inferring the numbers and stoichiometry of cohesin and HORMAD molecules in chromosome axis foci

Next, we set out to determine the numbers of the different cohesin and HORMAD proteins present in the individual axis foci that we observe cytologically.

First, we present evidence indicating that most individual REC-8 foci detected in fully spread nuclei represent single REC-8 cohesin complexes. We examined chromosome axes in fully spread meiotic chromosomes from worms expressing two differently-tagged versions of REC-8 (REC-8::3xFLAG and REC-8::GFP) in the same animal; no untagged REC-8 was expressed in these worms. If REC-8 foci represented cohorts of multiple REC-8 cohesin complexes, we would have expected to observe frequent colocalization of the two tags. Instead, visualization by immuno-fluorescence and SIM imaging revealed that both tagged proteins are integrated into the chromosome axes, but the signals corresponding to the two tags usually do not colocalize (FIG3A). Importantly, the same result was obtained when detection of FLAG and GFP was performed sequentially, in either order, indicating that lack of colocalization of the two tags was unlikely to have been caused by antibody interference (FIG3S1A).

**Figure 3.**
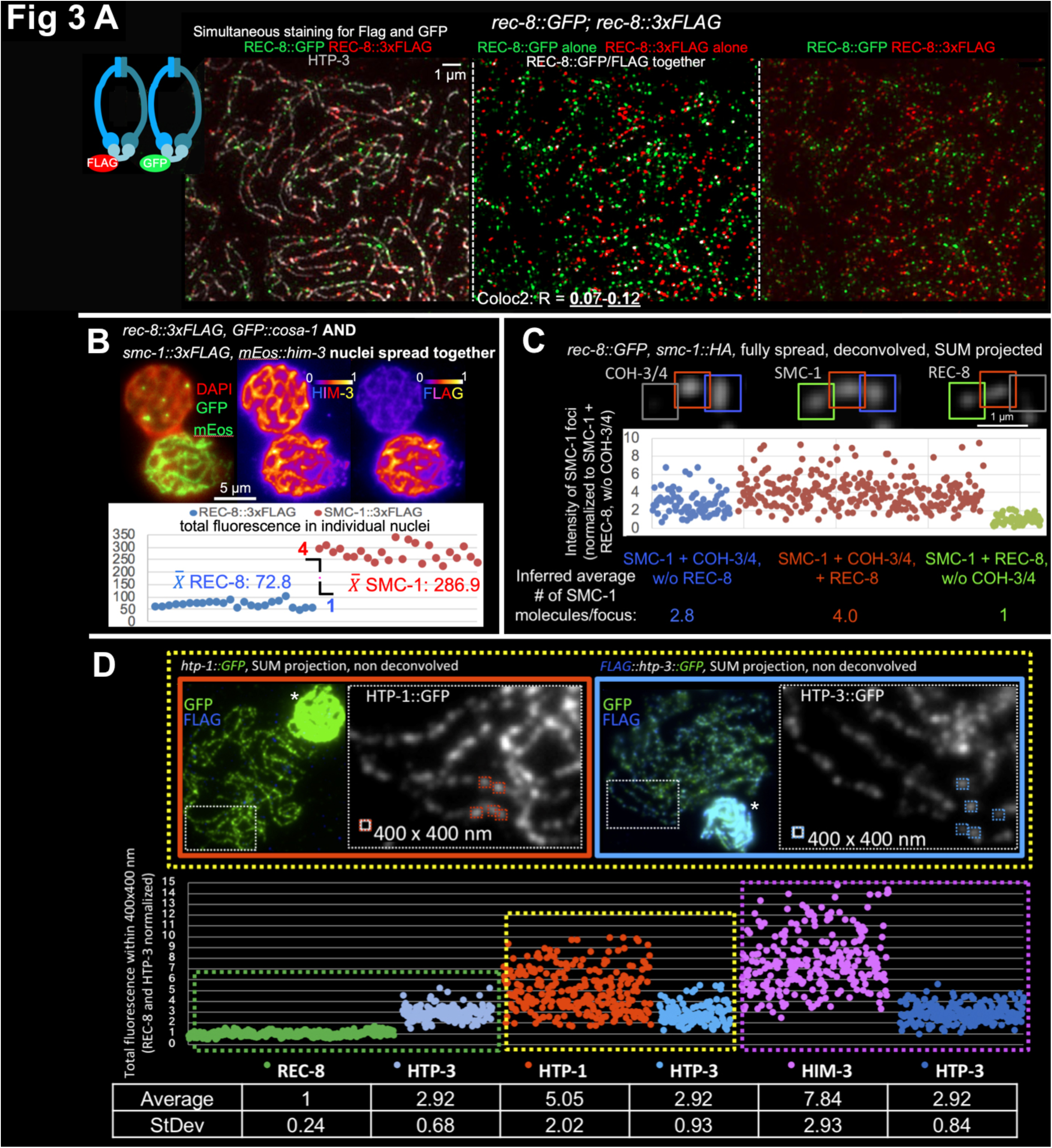
Quantitation of numbers of chromosome axis proteins present in individual axis foci. A) Sample SIM images of fully-spread nuclei from a worm expressing both REC-8::GFP and REC-8::3xFLAG, immuno-stained for HTP-3, FLAG and GFP. All primary antibodies (anti-FLAG, ant-GFP and anti-HTP-3) were applied at the same time. Middle panel was generated using ImageJ plugin “ColocThreshold”, with white indicating the infrequent pixels with significant FLAG and GFP colocalization (colocalizing pixel are in white, non-colocalizing pixel are in red and green); contrast and brightness were adjusted for display to improve visibility of the white pixels. Although GFP and FLAG signals both localize to chromosome axes (marked by HTP-3, left), they usually do not colocalize with each other. Coloc2 analysis using ImageJ confirmed the infrequent colocalization of FLAG and GFP by pixel intensity-based assessment (R(total) = 0.07-0.12; n=28 nuclei). B) Comparing the total amounts of chromatin-bound SMC-1 and REC-8. Gonads from worms expressing either SMC-1::3xFLAG and mEos::HIM-3 or REC-8::3xFLAG and GFP::COSA-1 were spread together on the same slides, immuno-stained for HIM-3 and FLAG, and imaged together using the same conditions. GFP::COSA-1 and mEos::HIM-3 served as internal markers to identify the genotypes of the nuclei being imaged; for the example nuclei depicted, similarities and difference in fluorescence intensities for HIM-3 and FLAG signals are illustrated using the indicated color scheme (generated using the Fire LUT from ImageJ). Graph indicates the total FLAG fluorescence measured in ROIs surrounding individual nuclei in SUM intensity projections; each data point represents a single nucleus. For each genotype, 6 late pachytene nuclei each from 4 different gonads were assayed. All values obtained are displayed, with no normalization applied. The ratio of the average total fluorescence values is 1 REC-8::3xFLAG : 3.94 SMC-1::3xFLAG. C) Inferred number of SMC-1 molecules in individual axis foci. The total fluorescence for SMC-1::HA, REC-8::GFP and COH-3/4 was measured in individual 800×800 nm ROIs surrounding individual foci in SUM intensity projections of fully spread nuclei; each data point represents the total fluorescence of SMC-1::HA in a single focus. As indicated in the cropped image on the top, SMC-1 foci were categorized as: localizing with COH-3/4, but not REC-8 (blue, left in the graph), localizing with both COH-3/4 and REC-8 (red, middle in the graph), or localizing with REC-8, but not COH-3/4 (green, right). All values obtained are plotted in the graph on the bottom. The Y-axis (representing arbitrary fluorescence units) is normalized to the mean fluorescence of SMC-1 foci that only localize with a REC-8 focus, but not with a COH-3/4 focus; this value is set to 1, as we consider REC-8 foci to represent single cohesin complexes. D) Sample images (top) and quantitative data (graph and table) for experiments measuring fluorescence intensities of individual foci in parallel for pairwise combinations of axis proteins. Nuclei from worms expressing 3xFLAG::HTP-3::GFP and nuclei from worms expressing REC-8::GFP, HTP-1::GFP or GFP::HIM-3 were spread together on the same slides, immuno-stained for GFP and FLAG, and imaged together using the same conditions. Presence or absence of FLAG staining (blue in the sample images) was used identify the genotype of the nuclei; asterisks indicate not unspread nuclei of the same genotype that were present in the same field. Total fluorescence was measured in 400×400 nm ROIs surrounding well-separated individual foci in SUM intensity projections; each data point represents a single focus. Different pairwise data sets were normalized to each other using the mean fluorescence of HTP-3::GFP foci as a normalization standard. The Y-axis (representing arbitrary fluorescence units) is normalized to mean REC-8::GFP = 1, as we consider REC-8 foci to represent single molecules.

We also employed an independent approach, using purified GFP protein as a single-molecule standard, to evaluate the numbers of REC-8::GFP molecules present in meiotic axis foci (FIG3S1B). Specifically, we spotted purified native GFP protein molecules on positively-charged slides, and we spread and fixed nuclei from worms that express REC-8::GFP (as the sole source of REC-8 protein) on top of them. We then compared the immunofluorescence signals associated with individual GFP protein molecules with those of REC-8::GFP axis foci, visualized either with monoclonal or polyclonal antibodies against GFP. We found that measured immunofluorescence signals for REC-8::GFP cohesin foci on chromosomes were similar to those measured for the individual GFP molecules (1.2:1).

Based on these analyses, we conclude that most of the individual REC-8 foci in the fully spread preparations each represents a single cohesin complex. We note that REC-8-containing cohesin complexes appear to be the only cohesins that connect sister chromatids during meiotic prophase in a classical cohesive manner, as COH-3/4 complexes not only bind to chromosomes after replication, but bind to individual chromatids and are insufficient to mediate sister chromatid cohesion in the absence of REC-8 (Severson and Meyer 2014; Crawley et al., 2016; Cahoon et al 2019; FIG2C). Thus, the fact that REC-8 (cohesive) cohesins usually occur locally as individual complexes is not consistent with the handcuff model for sister chromatid cohesion, which requires pairs of complexes to act together at any given site. Instead, our data strongly support the single ring hypothesis wherein sister chromatid cohesion is mediated locally by topological entrapment of two sister chromatids by a single cohesin ring.

Having established that single cytological REC-8 foci usually represent single REC-8 molecules, we set out to deduce the numbers and relative stoichiometry of other cohesin subunits and different HORMAD components associated with meiotic chromosome axes using quantitative analysis of wide-field immuno-fluorescence images.

We used two different independent approaches to infer the relative numbers of REC-8 and COH-3/4 containing cohesin complexes. First, we measured and compared the total amount of chromatin-bound SMC-1 (which is present in all cohesion complexes) and chromatin bound REC-8. Nuclei from gonads of two distinct genotypes, one expressing REC-8::FLAG (identified by GFP::COSA-1 expression) and the other expressing SMC-1::FLAG (identified by mEos::HIM-3 expression) were spread together on the same glass slide, immuno-stained for the Flag epitope, GFP (to detect GFP::COSA-1 and mEos::HIM-3) and HIM-3, and imaged using the same conditions (FIG3B). Whereas anti-HIM-3 and DAPI fluorescence were not distinguishable between the two genotypes (FIG3B and data not shown), the average total FLAG immuno-fluorescence signal for the nuclei expressing SMC-1::FLAG was 4 times that for the nuclei expressing REC-8::FLAG. Thus, we estimated there to be 4 times more chromatin/DNA bound SMC-1 than REC-8 molecules. We corroborated and extended this finding using an orthogonal approach in which we visualized and measured SMC-1 fluorescence signals (visualized with an HA tag) in 800nm × 800nm ROIs encompassing individual SMC-1 foci (FIG3C). SMC-1 fluorescence values were compared for three classes of foci, *i.e*., those associated with: a) COH-3/4 but not REC-8; b) both COH-3/4 and REC-8; c) REC-8 but not COH-3/4. The average SMC-1::HA fluorescence measured for “COH-3/4 only” foci was 2.8 x that measured for “REC-8 only” foci, and the average SMC-1::HA fluorescence for SMC-1 foci associated with both COH-3/4 and REC-8 was 4 x that for “REC-8 only” foci. Further, the total fluorescence measurements for COH-3/4 and for SMC-1 within the same foci are highly correlated (FIG3S1C), reinforcing the idea that REC-8 and COH-3/4 are the major, and possibly only, kleisins present in chromatin-bound cohesin complexes during meiosis (Severson et al., 2009). In summary, our two different quantification approaches together revealed that whereas individual REC-8 foci each represent one cohesion-mediating cohesin complex, COH-3/4 foci represent, on average, three axis-organizing cohesin complexes.

We also wished to determine the number and relative stoichiometry of different HORMAD components in the chromosome axis foci. To this end, we made pairwise comparisons of relative fluorescence intensities of foci for different axis components marked with the same tag, using HTP-3 as a standard for aligning the different comparisons. For each of these experiments, we isolated nuclei from two different genotypes (one expressing 3xFLAG::HTP-3::GFP and one expressing REC-8::GFP, GFP::HIM-3, or HTP-1::GFP), then mixed and spread them together on the same slide, and stained and imaged them using the same conditions. For the data in FIG3D, we measured fluorescence within 400×400 nm ROIs encompassing well-isolated individual foci and normalized our data to the mean fluorescence measured for REC-8 foci, as we consider these to represent single molecules. Using this approach, we estimated that a single cytological HORMAD focus contains, on average, 2.9 (+/-0.68) HTP-3 molecules, 5.1 (+/-2) HTP-1 molecules, and 7.8 (+/-2.9) HIM-3 molecules (FIG3D). The relative ratios of REC-8::GFP to HTP-3::GFP (1 : 2.9) and HTP-3::GFP to GFP::HIM-3 (1 : 2.7) calculated using this approach are in good agreement with those estimated using a different method that evaluates total fluorescence in partially spread nuclei (1 : 3.6 for REC-8::GFP to HTP-3::GFP and 1:3.2 for HTP-3::GFP to GFP::HIM-3) (FIG3S2A).

HORMAD proteins HTP-1 and HTP-2 are nearly identical, and although HTP-2 is largely dispensable for successful meiosis, they do have partially overlapping functions (Couteau and Zetka, 2005; Martinez-Perez et al., 2008; Martinez-Perez and Villeneuve, 2005). Thus, the value calculated for HTP-1::GFP is expected to be an underestimate of the total number of HTP-1/2 molecules associated with meiotic chromosome axes in nuclei that contain a mixture of tagged HTP-1 and untagged HTP-2. Indeed, we found that the ratio of HTP-1::GFP to HTP-3::GFP was significantly higher for nuclei that lacked HTP-2 (1.9 : 1) than for nuclei where HTP-2 was present (1.4 : 1) (FIG3S2B). Thus, we infer that a single HORMAD focus most likely contains an average of 6 HTP-1/2 molecules during WT prophase.

The wide ranges of fluorescence values measured for HTP-1 and HIM-3 foci likely reflect biological variation in the numbers of HTP-1 or HIM-3 molecules present in individual HORMAD foci, as we find that HTP-1/2 and HIM-3 protein levels in meiotic nuclei increase over course of meiotic prophase (FIG3S2C). In contrast, HTP-3 levels remain largely stable from meiotic entry through the end of pachytene (FIG3S2C). Overall, our data are in strong agreement with biochemical and structural data from a recent study that investigated interactions among *C. elegans* HORMAD proteins (Kim et al., 2014). This study found that the HIM-3 HORMA domain can bind to 4 of the 6 “closure” motifs in the C-terminal tail of HTP-3, and the HORMA domains of HTP-1/2 can bind to the remaining two motifs. Moreover, when the proteins are co-expressed in bacteria, HTP-3 and HIM-3 form complexes in a 1:2 or 1:3 ratio, while HTP-3 and HTP-1 do so in a 1:2 ratio. Thus, our data quantifying the numbers of foci per nucleus and their relative fluorescence are consistent with a model in which meiotic chromosome axes are assembled from interactions between individual cohorts of cohesin and HORMAD proteins in the following stoichiometry: [1 REC-8 cohesin] : [3 COH-3/4 cohesins] : [3 HTP-3 : 6-9 HIM-3 : 6 HTP-1/2], distributed along chromosomes with an average density of one such “module” for every 130-200 kbp based on the number of individual REC-8-containg cohesive cohesin complexes (FIG4A).

**Figure 4.**
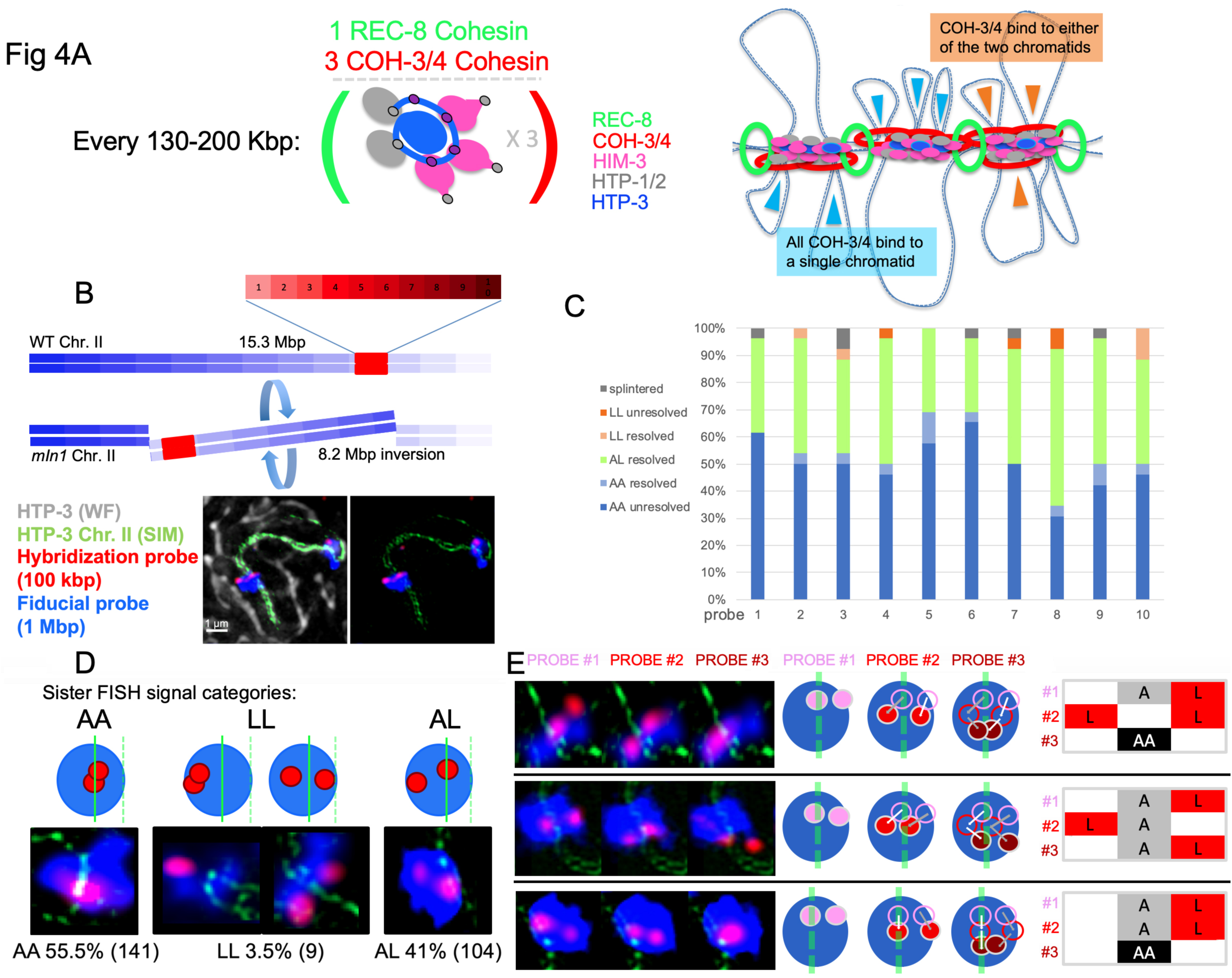
Model for chromosome axis organization and evidence for asymmetry between sister chromatid loops. A) LEFT: A quantitative model for chromosome axis foci composition, stoichiometry, and density derived from the data presented here and in (Kim et al., 2014). RIGHT: Model of chromosome axis organization based on the data presented; see text for description. B) Schematic illustrating the use of *mIn1*, a rearranged version of chromosome II harboring an 8.2 Mbp internal inversion, for immuno-FISH experiments assessing sister chromatid loop relationships. *mIn1* heterozygosity results in wide (>7 Mbp) separation between the FISH signals for the normal and rearranged homologs, thereby enabling assessment of sister chromatid signals. The red bar represents the 1 Mbp region covered by ten individual 100 kbp oligo paint FISH probes, which were detected sequentially. In the example image shown, a projected 3D-SIM image of the HTP-3 signals (green), corresponding to the individually cropped chromosome II bivalent from an *mIn1/+* heterozygous nucleus, is overlaid on the widefield image of HTP-3 (grey) for the whole nucleus. FISH signals from the 1 Mbp Fiducial probe are depicted in blue and the 100 kbp detection probe for the first hybridization step are depicted in red. Note that *mIn1* chromosome harbors an insertion within the inverted segment and therefore displays a longer axis than the WT chromosome *II*, with which it is engaging in synapsis. C) Stacked bar graph representing the frequency of occurrence of the indicated categories of sister-chromatid pair FISH signals for each of the ten consecutive individual 100 kbp oligo paint FISH probes assayed (1-10). (A: axis associated, L: loop) D) TOP: Schematic representation of categories of images observed for individual 100 kbp FISH probes. For each case, red circles indicate hybridization signals for an individual 100 kbp region, a blue oval indicates the hybridization signal for a fiducial probe corresponding to the entire 1 Mbp chromosome segment represented, and the solid green line indicates the chromosome axis corresponding to the hybridized chromosome. (The dashed green line indicates the axis corresponding to the synapsed partner chromosome, which is visible in some images but does not contain homologous DNA sequences at the same position owing to heterozygosity for a chromosomal inversion.) BOTTOM: Example images illustrating the individual categories of sister FISH signals schematized above, with the observed incidence of each type indicated below (n = 254; the 6/260 cases where FISH signals were splintered/diffuse were excluded from this analysis). E) Left, example images illustrating three consecutive sequential hybridization steps detected using consecutive 100 kbp probes. Center, cartoons illustrating the sequential hybridizations depicted. Right, corresponding schematic diagrams of the type used to codify and represent image galleries presented in FIG4S1.

### Model for chromosome axis assembly and implications for axis/loop organization

Based on our data and findings from previously-published literature (Severson et al., 2009; Severson and Meyer, 2014; Cahoon et al., 2019; Kim et al., 2014; Crawley et al., 2016), we propose the following model for the assembly and organization of meiotic chromosomes axes: Upon completion of pre-meiotic S-phase and entry into meiotic prophase, half of the REC-8-containing cohesin complexes, which were loaded during S-phase and hold sister chromatids together, are removed from chromatin by WAPL-1, resulting in connections between the two sister chromatids by a single REC-8 cohesin complex on average about every 130-200 kbp. Concurrently, COH-3/4-containing cohesin complexes bind and/or capture DNA of only one of the two sister chromatids, promoting the formation of loops in the intervals between the positions at which individual REC-8 complexes are maintaining cohesion between the sister chromatids. This organization is “locked in” by binding of a defined number of HTP-3 and other HORMADs, yielding an alternating occurrence of REC-8 cohesins and COH-3/4 cohesins along the resulting linear chromones axis (FIG4A).

When REC-8 is globally or locally absent, axis-promoting cohesin complexes (COH-3/4 in *C*. elegans or RAD21L in mouse) and HORMADs assemble functional chromosome axes and loops on single DNA molecules (FIG2C; Agostinho et al., 2016; Cahoon et al., 2019). Thus, we propose that on wild-type meiotic prophase chromosomes, COH-3/4-containing cohesin complexes also bind to individual chromatids between REC-8 mediated cohesion sites. In principle, the individual COH-3/4 complexes in each cytological focus (on average 3 complexes per cohort) could potentially bind to DNA from both sister chromatids (FIG4A, orange arrowheads), or alternatively, they could all bind in a coordinated manner to the same chromatid within any given interval between REC-8 cohesion sites (FIG4A, blue arrowheads). The first possibility could result in the formation of either symmetric or asymmetric loops, depending on the number of COH-3/4 complexes present in a given cohort. The latter possibility predicts that DNA loops that form on the two sister chromatids between two REC-8 cohesion complexes would be inherently asymmetric: DNA on one sister chromatid would form a single long loop that is not bound by COH-3/4, while the DNA on the other sister chromatid would be more closely associated to the chromosome axis in several shorter loops that are organized by a cohort of COH-3/4 complexes.

### Evidence for frequent asymmetry of sister chromatid loops

We used an iterative immuno-FISH (fluorescence *in situ* hybridization) approach (Mateo et al., 2019) to investigate the organization of the chromosome loops associated with pairs of sister chromatids, both relative to each other and in relationship to the chromosome axis. Specifically, we sequentially detected and imaged pools of oligo-paint FISH probes representing ten consecutive 100 kbp regions spanning a 1 Mbp region of chromosome II on chromosomes prepared using partial spreading conditions. To be able to distinguish FISH signals from sister chromatid pairs from signals derived from the homologous chromosome, we used worms heterozygous for an internal inversion of chromosome II (*mIn1*), which results in wide spacing between the FISH signals associated with the two homologs (FIG4B). Conventional deconvolution microscopy images of HTP-3 axis signals were acquired in conjunction with acquisition of images for each FISH detection step, enabling registration of images corresponding to consecutive genomic positions. In addition, a 3D-SIM image of the HTP-3 signal was acquired and used to enable assessment of the spatial relationships between FISH signals and the axis (see Methods).

Overall, most 100 kbp probe signals were detected either as a single unresolved focus associated with the chromosome axis (50%) or as two resolvable sister-chromatid foci (42%) with one or both sister signals clearly separated from the axis, *i.e*. in the loops (n = 260, FIG4C); the remaining cases consisted of unresolved single signals present in loops (1.5%), pairs of resolved sister signals that were both axis-associated (4%), or signals that were fragmented and/or noncompact (2.3%). Analysis of these immuno-FISH data yielded two important insights regarding axis/loop organization.

First, our data indicate that there is no locus within the assayed 1 Mbp chromosome segment that is consistently associated with the chromosome axis during the pachytene stage. For each of the ten probes, instances where one or both sister signals were clearly resolved from the axis were observed in at least 27% of hybridizations (FIG4C), indicating that none of the 100 kbp segments assayed contained DNA sequences that were consistently axis-associated. Thus, we conclude that the genomic sequences associated with the chromosome axis are variable between different meiocytes.

Second, analysis of this data set revealed strong evidence for asymmetry between sister chromatid loops (FIG4D-E). 100 kbp FISH signals that do not colocalize with the chromosome axis marker must represent DNA sequences that are located in chromosome loops (“L”). Conversely, FISH signals that correspond to axis-associated DNA, or DNA in associated loops that are smaller than 100 kbp in size, will colocalize with the axis marker (“A”). If sister chromatid loops were consistently symmetric, we would expect that for any given 100 kbp DNA segment, sister FISH signals would typically either both be axis-associated (“AA”) or both be in loops (“LL”), and cases where one sister signal was axis-associated and the other was in a loop (“AL”) would be rare. Our observed data clearly depart from this expectation (p< 0.00001), as 41% of sister FISH signal pairs were of the AL category and only 3.5% were of the LL category, with the rest being of the AA category (55.5%, n = 254 loci assayed, FIG4D). At face value, the observed distribution of AA, AL and LL signal pairs would appear to be compatible with that expected for independent behavior of the two sister chromatids (p = 0.1443). However, a different picture emerges when the expected behavior at sites of REC-8-mediated sister chromatid cohesion is taken into consideration. Using the conservative estimate of an average density of one REC-8 cohesin complex every 200 kbp (FIG1), half of the 100 kbp probes in any given hybridization sequence would be expected to include a REC-8-mediated cohesion site and thus would yield an “obligate” unresolved axis-associated (AA) signal in the immuno-FISH analysis. If these obligate AA sites are removed from consideration when evaluating behavior of sister chromatid FISH signals, the new distribution of the remaining 127 signal pairs is 11% AA, 82% AL and 7% LL, i.e., AL signal pairs (in which sister chromatids exhibit different behavior) represent the preponderance of cases. This clearly differs from expectations for independent behavior of sister chromatids (p < 0.00001), and instead implies a strong propensity for sister chromatids to be asymmetric. Consistent with this interpretation, when we analyzed 40 sequences of consecutive FISH detection steps for individual chromosomes and considered the possible paths of loops that were >100 kbp in size (see FIG4S1 and legend), we observed only a single case where loops of the same size occurred at the same position on both chromatids (FIG4E, FIG4S1 and data not shown).

As spreading of chromosomes and Z-projection of images might potentially introduce artifacts, we carried out a separate assay using a 200 kbp FISH probe in minimally disrupted tissue analyzed in 3D (FIG4S2A). This approach corroborated our finding that the majority of chromosome loops are asymmetric. Specifically, we found that the majority of sister FISH signal pairs were in the AL class (57%), with only 5% in the LL class and the remainder (38%) in the AA category (n = 94 loci assayed, FIG4S2B). Thus in summary, we conclude that the loops present on pairs of sister chromatids are frequently distinct from each other in size and/or position, and consequently, in genomic content. Indeed, our data are consistent with a model in which sister chromatid loops are inherently asymmetric.

Asymmetry between sister chromatid loops may be a widespread feature of meiotic prophase, as loop asymmetry was observed previously for giant loops of diplotene-stage lampbrush chromosomes of the newt (Fairchild and Macgregor, 1994). Our observation of sister loop asymmetry during an earlier stage of meiotic prophase can potentially help to explain some important properties of the meiotic recombination program. Since the DSBs that serve as the initiating events of meiotic recombination are thought to be generated in loop regions that are tethered to the chromosome axis (Blat et al., 2002; Panizza et al., 2011), such asymmetry between sister-chromatid loops could potentially help to prevent DSBs from occurring at the same site on both sister chromatids during the same meiosis. Furthermore, the proposed axis-loop organization could also contribute to the inter-homolog repair bias of meiotic recombination (Haber et al., 1984; Jackson and Fink, 1985; Schwacha and Kleckner, 1994), as an intrinsic feature of the proposed configuration is that sister chromatids would be unlikely to share the same loop domains and thus would not be favored as DNA repair partners.

### Concluding remarks

The approaches employed in this work have enabled us to deduce the numbers, relative stoichiometry and spatial distribution of multiple proteins that are central to the functional organization of chromosomes during meiosis. Further, our data regarding the number of cohesin foci and the stoichiometry of proteins within these foci can be used to calculate the density of cohesin complexes along *C. elegans* meiotic prophase chromosomes. Based on our evidence that most REC-8 foci represent single cohesin complexes and that COH-3/4 foci represent, on average, 3 cohesin complexes, we can infer that there are approximately 20 - 30 cohesin complexes per Mbp of chromosome length when each pair of sister chromatids is considered as a single conjoined entity. When considering the total amount of DNA in each sister chromatid pair, this translates to a density of 10 - 15 cohesin complexes per Mbp of DNA. These measurements of cohesin complex density on *C. elegans* meiotic chromosomes are in remarkably close agreement with recent independent estimates of cohesin density deduced for mouse ES cells based on in-gel fluorescence measurements and fluorescence correlation spectroscopy (FCS)-calibrated imaging (5.3 complexes/Mb; (Cattoglio et al., 2019)) and for HeLa cells based on quantitative mass spectrometry and FCS-calibrated imaging (8.5-17 complexes/Mb; (Holzmann et al., 2019)), supporting both the validity of our experimental approach and the broad relevance of our findings to the field of chromosome biology. Thus, we anticipate that other features revealed in the current work, such as mediation of cohesion by individual cohesin complexes, and spatial alternation between cohesion-mediating cohesin complexes and putative axis/loop-organizing cohesin complexes, may reflect generalizable principles and properties of chromosome organization that operate in other contexts. Finally, we note that the approaches that we devised here to quantify numbers and/or relative stoichiometry of chromosome axis components should be widely applicable for analyzing many other cellular structures and biological processes, enabling quantitative estimates of molecular components that can inform and constrain models regarding how such structures and processes assemble and function.

## MATERIALS AND METHODS

### *C. elegans* culture conditions

Worms were grown on *E. coli* (OP50) seeded NG agar plates at 20°C according to the standard method (Stiernagle, 2006). For experiments, worms were either selected as homozygous L4 larvae (from heterozygous strains maintained using balancer chromosomes) or were harvested as staged L1 larvae following bleaching of gravid adults (for strains maintained as homozygotes or for *mIn1/+* heterozygotes) according to: (Stiernagle, 2006) and were analyzed 24-36 hours post L4 stage. For experiments using two tagged versions of REC-8, ATGSi23 (*rec-8::GFP; rec-8 (null)*) hermaphrodites were mated with CA1481 (*rec-8::3xFLAG*) males, and leptotene/zygotene nuclei of F1 heterozygous hermaphrodites, which expressed both REC-8 tagged proteins (and no untagged REC-8), were analyzed 36 hours post L4 stage.

*C. elegans* strains used in this study:

- N2
- ATGSi23: *fqSi23[Prec-8::rec-8:::GFP::rec-8 3’UTR + cb-unc-119(+)] II; rec-8(ok978) IV*
- CA1481: *mels8[Ppie-1::GFP::cosa-1 + cb-unc-119(+)] II; rec-8(ie35[rec-8::3xFLAG]) IV*
- TY4986: *htp-3(y428) ccls4251 I/hT2 (I, III)*
- ATG98: *wapl-1(tm1814) IV/nt1[qIs51] (IV,V)*
- TY5120: *+/nT1 IV; coh-4(tm1857) coh-3(gk112) V/nT1[qls51] (IV;V)*
- VC666: *rec-8(ok978) IV/nT1[qls51] (IV;V)*
- AV1079: *meEx001; rec-8(ok978)/nT1[qls51] (IV;V)*
- EG699: *ttTi5605 II; unc-119 (ed3); oxEx1578*
- CA1282: *him-3(ie114[gfp::him-3]) IV*
- CA1437: *htp-3(tm3655) I; ieSi62[Phtp-3::3×Flag::htp-3::GFP::htp-3 3′UTR + unc-119(+)] II; him-3(ie33[mEos2::him-3]) IV*
- ATGSi43: *fqSi37[htp-1::GFP cb-unc-119(+)] II; htp-1(gk174) IV*
- AV1080: *fqSi37[htp-1::GFP cb-unc-119(+)] II; htp-1(gk174) htp-2(tm2543) IV*
- AV1088: *smc-1(ie39[smc-1::intHA]) I; fqSi23[Prec-8::rec-8:::GFP::rec-8 3’UTR + cb-unc-119(+)] II; rec-8(ok978) IV*
- VC1474: *top-2(ok1930)/mIn1[mls14 dpy-10(e128)]* II
- CA1432: *smc-1(ie36[smc-1::3xFlag]) I; mels8[Ppie-1::GFP::cosa-1 + cb-unc-119(+)] II; him-3(ie33[mEos2::him-3]) IV*

#### Microinjection of *C. elegans*

Microinjection was performed using a FemtoJet 4i injector system (Eppendorf) and an Axiovert 10 microscope (Zeiss). Injection mixes contained a total of 100ng DNA in distilled water. The injection mix for *meEx001* contained the dominant selection plasmids pGH8 (*prab-3::mCherry::unc-54_3’UTR*, 2.5ng) and pBR186 (modified from pDD282, includes *sqt-1p::sqt-1(e1350):: sqt-1_3’UTR _hsp16::Cre::tbb-2_3’UTR_rps-11p::HygR::unc-54_3’UTR*, 2.5ng) in addition to 95 ng *E. coli* (OP50) DNA, isolated by standard protocols, sonicated and size-selected for fragments of 1-5 kbp by agarose gel purification.

#### Full spreading and staining of nuclei

Worms were staged by bleaching, grown at 20°C and harvested 24-36h post L4 stage (pelleted by gravitation), washed 3 times in 0.35x nuclear purification buffer (NPB; 3.5 mM HEPES pH 7.5, 14 mL NaCl, 31.5 mM KCl, 0.07 mM EDTA, 0.175 mM EGTA, 0.07 mM DTT, 0.035% Triton X-100). The resulting worm pellet was mixed with 1 volume 0.35x NBP and frozen as worm beads by dropping 50 μl drops of NBP-worm slurry into liquid nitrogen. 2-10 beads were ground for 10 seconds using a liquid nitrogen cooled mortar and pestle. The ground tissue was transferred into a 50 ml Falcon with a kitchen spoon, thawed and pipetted up and down several times to release nuclei. 5 μl of this suspension was applied on an EtOH-washed 22×40mm coverslip. 50μl of spreading solution (see below) was added and nuclei were immediately distributed over the whole coverslip using a pipette tip. Coverslips were left to dry at room temperature (approximately 1 hour) and post-dried for two more hours at 37°C, washed for 20 minutes in methanol at -20°C and rehydrated by washing 3 times for 5 minutes in PBS-T. A 20-minute blocking in 1% w/v BSA in PBS-T at room temperature was followed by overnight incubation with primary antibodies at 4°C (antibodies diluted in: 1% w/v BSA in PBS-T supplied with 0.05% w/v NaN_3_). Coverslips were washed 3 times for 5 minutes in PBS-T before secondary antibody incubation for 2 hours at room temperature. After PBS-T washes, the nuclei were immersed in Vectashield (Vector) and the coverslip was mounted on a slide and sealed with nail polish. Spreading solution: (for one coverslip, 50μl): 32μl of Fixative (4% w/v Paraformaldehyde and 3.2–3.6% w/v Sucrose in water), 16μl of Lipsol solution (1% v/v/ Lipsol in water), 2μl of Sarcosyl solution (1% w/v of Sarcosyl in water).

#### Partial spreading and staining of nuclei

Partial spreading of *C. elegans* gonads was performed as in (Pattabiraman et al., 2017). The gonads of 20–100 adult worms were dissected in 5μl dissection solution (see below) on an EtOH-washed 22×40mm coverslip. 50μl of spreading solution (see above) was added and gonads were immediately distributed over the whole coverslip using a pipette tip. Coverslips were left to dry at room temperature (approximately 1 hour) and post-dried for two more hours at 37°C, washed for 20 minutes in methanol at -20°C and rehydrated by washing 3 times for 5 minutes in PBS-T. A 20-minute blocking in 1% w/v BSA in PBS-T at room temperature was followed by overnight incubation with primary antibodies at 4°C (antibodies diluted in: 1% w/v BSA in PBS-T supplied with 0.05% w/v NaN_3_). Coverslips were washed 3 times for 5 minutes in PBS-T before secondary antibody incubation for 2 hours at room temperature. After PBS-T washes, the nuclei were immersed in Vectashield (Vector) and the coverslip was mounted on a slide and sealed with nail polish. Dissection solution: For Figures 1A (second nucleus on the top left), 1S1B, 1S2A, 2A, 2S1A, 3S2A, 4 and 4S1 gonads were dissected in 10-30% v/v Hank’s Balanced Salt Solution (HBSS, Life Technology, 24020-117) with 0.1% v/v Tween-20. For Figures 1S2B, 2C, 2S1C, and 4S2 gonads were dissected in 50-85% v/v Hank’s Balanced Salt Solution (HBSS, Life Technology, 24020-117) with 0.1% v/v Tween-20.

#### Antibodies used in this study

The following primary antibodies used: Chicken anti-HTP-3 (1:500) (MacQueen et al., 2005), chicken anti-GFP (1:2000) (Abcam), mouse anti-GFP (1:500) (Roche), rabbit anti-GFP (1:500) (Yokoo et al., 2012), mouse anti-HA (1:1000) (Covance/Biolegend), guinea pig anti-SYP-1 (1:200) (MacQueen et al., 2002), mouse anti-H3K9me2 (1:500) (Abcam), rabbit anti-SMC-1 (1:200) (Chan et al., 2003), rabbit anti-COH-3/4 (1:5000) (Severson and Meyer, 2014), rabbit anti-REC-8 (1:10000) (Novus Biologicals), rabbit anti-HIM-3 (1:200) (Zetka et al., 1999), rabbit anti-HTP-1/2 (1:500) (Martinez-Perez et al., 2008) and mouse anti-FLAG (1:200) (Sigma).

Secondary antibodies conjugated to Alexa dyes 405, 488, 555 or 647, obtained from Molecular Probes, were used at 1:500 dilution (Alexa 488 and 555), 1:200 (Alexa 647) and 1:100 (Alexa 405). In cases where antibodies raised in mouse and guinea pig were used on the same sample, we used highly cross-absorbed goat anti-mouse secondary antibodies, obtained from Biotium (conjugated to CF488, or CF555 respectively) for secondary detection of the mouse primary antibody in order to avoid cross-reaction against antibodies raised in guinea pig.

#### Imaging

Imaging, deconvolution and 3D-SIM reconstruction were performed as in (Pattabiraman et al., 2017). Wide field (WF) images were obtained as 200 nm spaced Z-stacks, using a 100x NA 1.40 objective on a DeltaVison OMX Blaze microscopy system, deconvolved and corrected for registration using SoftWoRx. Subsequently, gonads were assembled using the “Grid/Collection” plugin (Preibisch et al., 2009) in ImageJ. For display, pictures were projected using maximum intensity projection in ImageJ. 3D-Structured Illumination microscopy images were obtained as 125 nm spaced Z-stacks, using a 100x NA 1.40 objective on a DeltaVison OMX Blaze microscopy system, 3D-reconstructed and corrected for registration using SoftWoRx. For display, images were projected using maximum intensity projection in ImageJ or SoftWoRx. For display in figures, contrast and brightness were adjusted in individual color channels using ImageJ.

#### Manual quantification of axis foci

Axis foci were counted manually on 32-bit Z-projected SIM images. For this analysis, foci are defined as fluorescence signals that 1) display individual maxima in non-overlapping ROIs ≥ 3×3 pixels in size and 2) were at least five times brighter than background average (with most ranging from 10-100 times brighter than background).

#### Quantitation of relative levels of chromosome axis proteins present in individual axis foci

Nuclei from worms expressing 3xFLAG::HTP-3::GFP and nuclei from worms expressing REC-8::GFP, HTP-1::GFP or GFP::HIM-3 were isolated in parallel, mixed, and spread together on the same slides. GFP was labeled using primary and secondary antibodies prior to primary and secondary detection of the FLAG epitope to identify the nuclei expressing 3xFLAG::HTP-3::GFP, and nuclei on the same slide were imaged in parallel using the same conditions. SUM intensity projections were generated from non-deconvolved 32-bit images, and total fluorescence was measured in 400×400 nm ROIs surrounding well-separated individual foci. Fluorescence measurements were made for 25-70 well-separated foci per nucleus (4-9 nuclei for each axis component); each data point represents a single focus. Different pairwise data sets were normalized to each other using the mean fluorescence of HTP-3::GFP foci as a normalization standard. The Y-axis (representing arbitrary fluorescence units) is normalized to mean REC-8::GFP = 1, as we consider REC-8 foci to represent single molecules.

#### Assessing localization of REC-8 and COH-3/4 on straightened axes

Axis segments that did not twist around the homologous partner locally and were not spread on top of each other in maximum intensity Z-projected SIM images were manually traced and straightened using the ImageJ ‘‘Segmented Line Tool” and “Selection straighten” operation as a 10 pixel-wide band. Signal peaks in individual channels were manually identified and marked as a single pixel along the band. The individual channels were analyzed separately, and the outputs were overlaid prior to manual readout of the spatial relationships of peak positions, resulting in the graphs presented in FIG3.

#### Colocalization analysis using Coloc2 and Colocalization Threshold

Colocalization was assessed by evaluating the correlation of pixel intensities over space using the Coloc2 plugin of ImageJ. R values around 0 indicated lack of colocalization while an R value of 1 would indicate 100% colocation of signals in two channels, with their intensities completely correlated. 32-bit images were cropped and transformed into 8-bit single channels images. Background was subtracted in ImageJ using the sliding paraboloid background subtraction operation before running the Coloc2 or the Colocalization Threshold analyses.

#### Measuring total fluorescence of single axis-component foci

Nuclei from animals expressing 3xFLAG::HTP-3::GFP and nuclei from animals expressing REC-8::GFP, HTP-1::GFP or GFP::HIM-3 were isolated, mixed and spread on the same slides in a pairwise manner. GFP was visualized with a primary antibody raised in chicken followed by secondary detection with an Alexa488-conjugated anti-chicken antibody prior to primary and secondary detection of the FLAG epitope to distinguish between nuclei expressing 3xFLAG::HTP-3::GFP and REC-8::GFP, HTP-1::GFP or GFP::HIM-3; this approach also served to avoid potential (but not observed) antibody exclusion artifacts when co-detecting FLAG and GFP on the same molecule. Nuclei on the same slide were imaged using the same conditions; SUM intensity projections were generated from non-deconvolved 32-bit images, and total fluorescence was measured in 400×400 nm ROIs surrounding well-separated individual foci. Fluorescence measurements were made for 25-70 well-separated foci per nucleus (4-9 nuclei for each axis component); all data point measured are presented in the graphs.

#### 3D-tracing of chromosomes

32-bit SIM images obtained from high salt spreads were transformed first into RBG color images and then into monochrome 8-bit images. In such images, individual chromosomes were traced using the “Simple Neurite Tracer” plug-in of ImageJ (Longair et al., 2011). For display and straightening of individual chromosomes, each individual chromosome path was exported as an 8-bit two-color mask with the “Fill Out” function of the “Simple Neurite Tracer”. This mask was used to crop the individual chromosome in 3D from an RGB stack. These pictures were projected in Z and straightened using ImageJ ‘‘Segmented Line Tool” and the “Selection straighten” operation.

#### Comparing fluorescence of purified GFP protein and REC-8::GFP foci

Purified His-tagged monomeric GFP protein (a gift from P. Jackson) was diluted to 0.3 mM concentration in PBS, applied to positively charged slides (“Superfrost Plus”, Fisher Scientific) and incubated for 30 minutes in a moist chamber.

Most of the liquid was aspirated off, and immediately afterward, nuclei isolated from worms expressing REC-8::GFP (in a *rec-8* deletion background; ATGSi23) were spread on top using the “Full spreading protocol” with no alterations.

GFP was visualized using rabbit polyclonal or mouse monoclonal anti-GFP primary antibodies and Alexa 647-labelled secondary antibodies (Molecular Probes/Invitrogen) directed against rabbit or mouse, respectively. In a sequential step, the chromosome axis was visualized by anti-HTP-3 and Alexa 555-labelled anti-chicken secondary (Molecular Probes/Invitrogen). Non-deconvolved wide-field images were SUM-intensity projected. Fluorescence signals were measured for 4×4 pixel ROIs encompassing individual well-spaced REC-8::GFP foci or purified GFP protein foci from the same image. Background in the image was measured in same-sized ROIs, averaged and subtracted. All obtained values are presented.

#### Whole-nucleus comparisons of anti-FLAG fluorescence

Gonads from worms expressing GFP::COSA-1 and REC-8::3xFLAG were dissected and mixed with gonads from worm expressing 3xFLAG::HTP-3::GFP and spread on the same slide using partial spreading. To avoid antibody exclusion artifacts, only the FLAG tag was detected by anti-FLAG immuno-staining, and GFP was detected using native GFP fluorescence. Gonads were additionally stained with DAPI. Genotypes were distinguished by GFP::COSA-1 or HTP-3::GFP expression, respectively. Z-stacks were acquired using the same wide-field conditions and SUM-intensity projected without prior deconvolution. ROIs were drawn encircling late pachytene nuclei from three gonads for each genotype (20 nuclei per gonad). Total FLAG and DAPI fluorescence within each ROI were recorded; values presented in the graph are ratios between FLAG and DAPI fluorescence (to normalize for differences in numbers of Z planes occupied by the nuclei in the SUM projection and differences in ROI sizes). Values were further normalized to: average value of REC-8::GFP = 1. All obtained values are presented.

#### Design and generation of Oligopaint FISH probes

Probes were designed to target a 1 Mbp region on chromosome II, between coordinates 11,500,001-12,500,001, as in (Mateo et al 2019). The 1 Mbp region was divided into 100 10 kbp segments, and 100 probes were targeted to hybridize to each 10 kbp segment. Each probe consisted of: 1) a unique non-overlapping 40-nt sequence corresponding to genomic DNA, 2) a 20-nt barcode sequence identifying each 10 kbp segment, 3) a 20-nt fiducial probe binding sequence (catcaacgccacgatcagct), and 4) a unique pair of index sequences at the 5’ (ACGTCCGCCGCATCTACGAG) and 3’ (CTGAACGGCGCACGTGTCTT) ends that allowed this probe set to be specifically amplified from a larger oligopool ordered from CustomArray. The probe set was synthesized from the oligopool as in (Mateo et al 2019), using forward primer (ACGTCCGCCGCATCTACGAG) and reverse primer (AAGACACGTGCGCCGTTCAG). Oligopaint probe sequences are provided in Supplementary Data File 1; read-out probe sequences and strand-displacement oligo sequences are provided in Supplementary Data Tables 1 and 2.

#### Sequential immuno-FISH experiments

Gonads from animals heterozygous for *mIn1* were dissected and spread on a coverslip using partial spreading conditions. After the sample was dried, it was incubated in -20C methanol for 20 minutes. Then, the sample was rehydrated by washing in PBST for 5 minutes each. Next, the sample was incubated in 0.1M HCl for 5 minutes and washed in PBST 3 times for 5 minutes each. The samples were then incubated for 5 minutes each in increasing concentrations of formamide in 2x SSCT: 5%, 10%, 25%, and 50%. The sample was then incubated in pre-warmed 42C solution of 50% formamide in 2x SSCT for 1 hour. 2μl of Oligopaint probe (1000ng/μl in dH2O) were diluted in 30μl of hybridization solution (10% Dextran Sulfate solution, 20X SSC, fdH2O, 10 μl 10% Tween for 1 ml of hybridization solution) for each coverslip. After 1 hour incubation, slides were taken out of the 50% formamide solution, wiped, and incubated in 95% ethanol for 5 minutes. Then, the probe hybridization solution was applied to the sample with a slide and the sample was denatured for 10 minutes at 90C on a heat block. After denaturing, the sample was incubated with the probe hybridization solution at 42C overnight. The next day, samples were washed 2 times in 42C 50% formamide in 2x SSCT for 30 minutes each and the slide was removed from the coverslip. Then, the sample was incubated for 5 minutes each in solutions with decreasing concentrations of formamide in 2x SSCT: 25%, 10%, and 5%. Samples were then washed 2 times for 10 minutes each in 2x SSCT and post fixed in 2% formaldehyde and 8% glutaraldehyde in 1xPBS followed by three 5 minute washes in PBST. The sample was blocked in 1% w/v BSA in PBS-T at room temperature and further incubated overnight with a primary antibody against HTP-3 at room temperature (antibodies diluted in: 1 % w/v BSA in PBS-T supplied with 0.05% w/v NaN_3_). Coverslips were washed 3 times for 5 minutes in PBS-T and blocked again in 1% w/v BSA in PBS-T. Coverslips were incubated with a secondary antibody conjugated to Alexa 488 for 2 hours at room temperature, and subsequently washed 3 times for 5 minutes in PBS-T, and samples were covered in Vectashield. A homebuilt microfluidic chamber was mounted onto to the coverslip with silicon, and this assembly was mounted onto a homebuilt holder for imaging using the OMX Blaze system.

In order to visualize the hybridization probes in 10 sequential 100 kbp steps, readout oligos (which hybridize to barcode sequences specific to each 10 kbp in the oligopaint library) were pooled in groups of 10 for each hybridization step, and readout probes were labeled indirectly via hybridization with Cy5-marked ‘imaging’ oligos (Cy5-5p-TGGGACGGTTCCAATCGGATC). A fiducial probe, which labels the entire 1 Mbp region, was marked by Cy3. Sequential hybridization of readout and imaging oligos and probe displacement steps were performed as in (Mateo et al., 2019). At each hybridization step, widefield image stacks of mid-pachytene nuclei (in 50% Vectashield in 1xSCC) were acquired for all three channels. A 3D-SIM image of the HTP-3 and fiducial signals was acquired after the fifth round of hybridization. Widefield image stacks were deconvolved and corrected for drift in 3D with ImageJ Plugin: “Correct 3D drift” using first the fiducial probe signal and then HTP-3 as a reference channel. Images were SUM-intensity projected and doubled in size to fit the 3D-SIM image size. Brightness differences between individual hybridization/imaging steps were corrected with ImageJ Bleach Correction (using “Histogram matching”) in all three channels. Chromosome axes for which fiducial probes were unambiguously assignable were cropped individually from 3D-SIM images and overlaid with widefield HTP-3 signals in the SUM projected pictures. When axes were imaged from a frontal view *i.e*. when two roughly parallel axis signals could be detected, the fiducial probe signal was inferred to correspond to DNA associated with the closer of the two axis signals. Only cases where an unambiguous assignment could be made were analyzed. Individual signals for each hybridization step that fully or partially overlapped with HTP-3 signals were considered “axis-associated”. Signals not colocalizing with HTP-3 were considered to be in “loops”. Brightness and contrast were adjusted for display.

## ACKNOWLEDGMENTS

We are grateful to A. Dernburg, B. Meyer, A. Severson and E. Martinez-Perez for antibodies and strains. We thank J. Mulholland, K. Lee, A. Kahn and C. Akerib for technical assistance and discussions and A. MacQueen, C. Jacovetti and C. Akerib for comments on early versions of the manuscript. This work was funded by an American Cancer Society Research Professor Award (RP-15-209-01-DDC) and NIH grants R01GM53804 and R35GM126964 to AMV, a FWF Erwin Schrödinger Fellowship (J-3676) to AW and grant 1S10OD01227601 from the NCRR to the Stanford Cell Science Imaging Facility.

## AUTHOR CONTRIBUTIONS

Investigation: AW and KY. Conceptualization, Methodology: AW. Formal Analysis, Funding Acquisition: AMV and AW. Writing - original draft: AW. Writing - review and editing, Resources: AMV, AW, KY, BR, SK and AB.

## DECLARATION OF INTRESTS

The authors declare no competing interests.

## Supplementary Data Files for

**Supplementary Data File 1:**

This file contains Oligopaint Probe Sequences as a fasta file.

**Supplementary Data Table 1:**

This file contains Read-out probe sequences as an Excel file.

**Supplementary Data Table 2:**

This file contains Strand-displacement oligo sequences as an Excel file.

**Figure 1S1.**
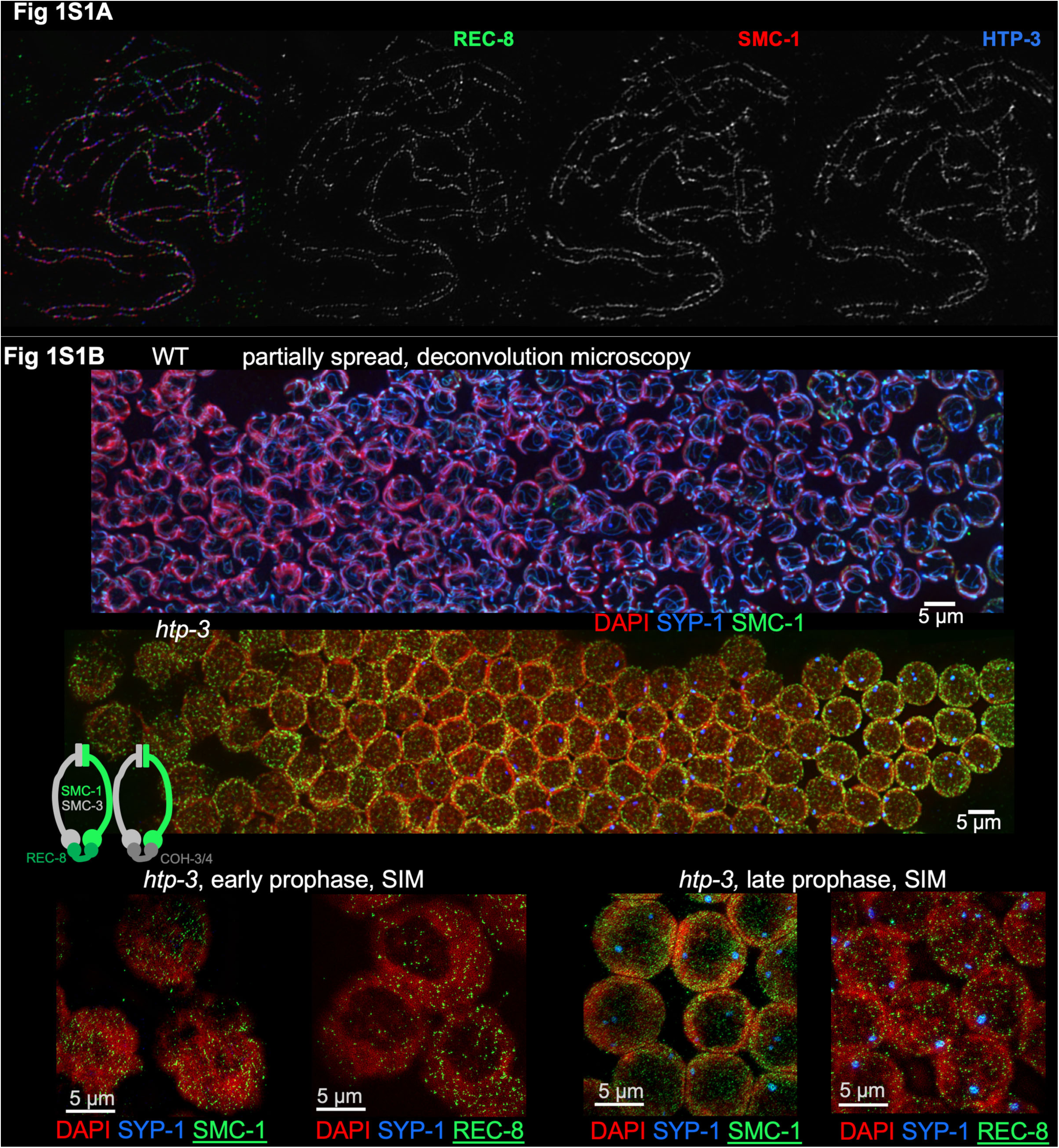
Requirement for HTP-3 in organizing cohesins into axial arrays. A) SIM image of a fully spread nucleus stained for REC-8::GFP, SMC-1 and HTP-3. B) Images of cohesin components SMC-1 and REC-8 and SC central region component SYP-1 in partially spread nuclei from *htp-3* mutant gonads, visualized by wide-field deconvolution microscopy. SMC-1 and REC-8 are not detected as linear arrays of foci or as continuous axes in either early or late prophase nuclei (indicated by the presence of 1-3 SYP-1 aggregates).

**Figure 1S2.**
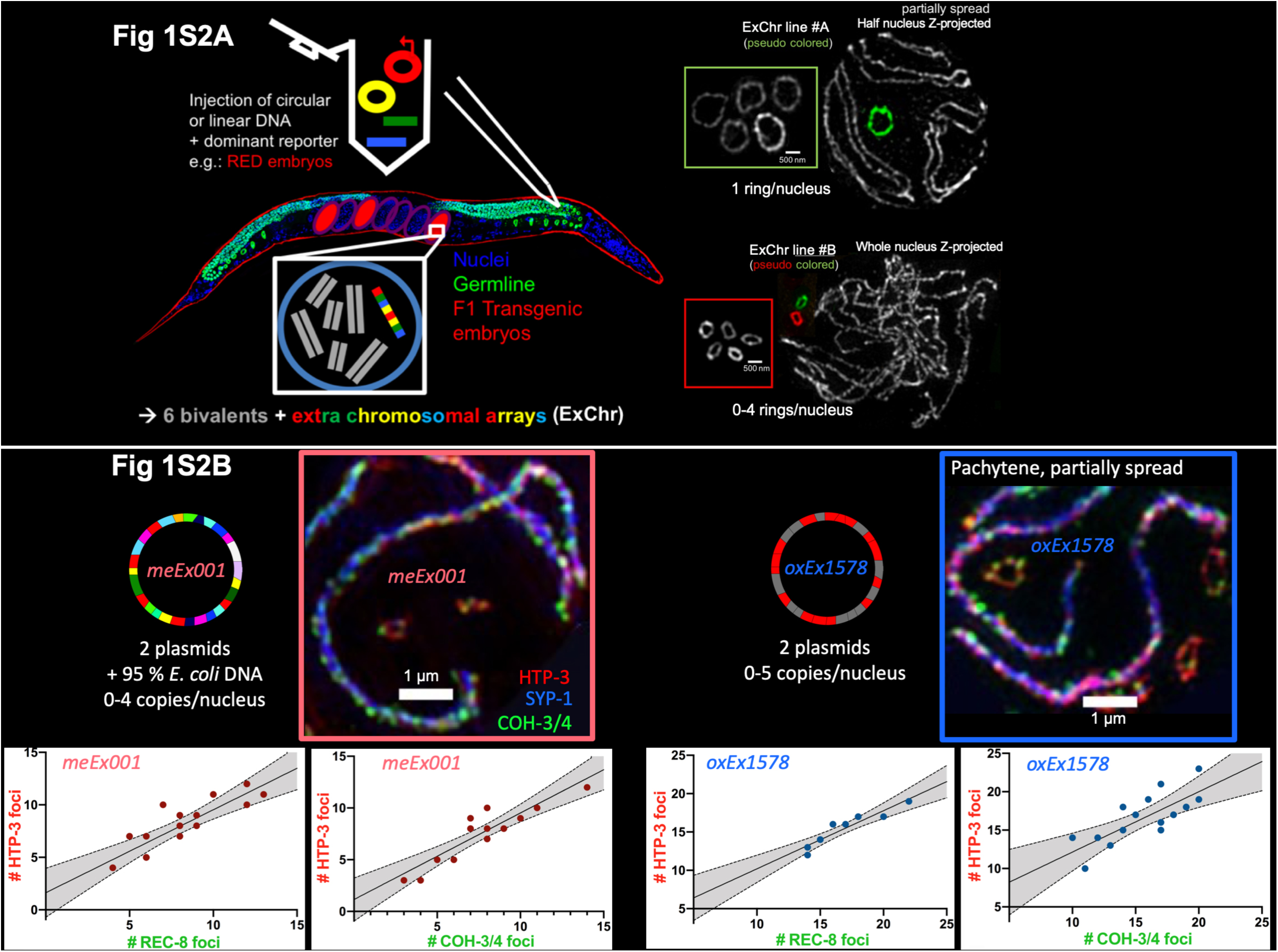
Structure and features of ExChrs. A) Circular structure of ExChrs. Left, schematic illustrating that multiple circular and/or linear DNA molecules co-injected into the *C. elegans* germ line co-assemble into extra-chromosomal arrays that can be inherited as mini-chromosomes. Right, SIM images of HTP-3-marked chromosome axes in spread nuclei carrying ExChrs, which appear as rings. Insets show multiple rings cropped from different nuclei from the same strain; the fact that rings from a given strain are similar in size indicates that once formed, ExChrs tend to be stable in size. The rings depicted are two different examples from 21 independent lines generated using the same selection scheme [*cb-unc-119(+)* in the *unc-119(ed3)* mutant], either alone or together with 10-90% *E. coli* genomic DNA (linearized by sonication and size selected to 1-5 kbp by gel extraction) in the injection mix. The transmission rates of the Unc+ trait in self-fertilizing hermaphrodites varied widely, ranging from 10-95%, and did not correlate with the percentage of *E. coli* DNA present in the injection mix. In 18 of the 21 lines, ExChrs could be detected as additional HTP-3-positive structures in meiotic prophase nuclei, and in 16 of these 18 cases, the ExChrs were reliably resolvable as rings by SIM in partially spread gonads; as the two ExChrs that were not verified as rings were the smallest of the 18, we speculate that they may also be rings, but their small size prevented their ring structure from being resolved microscopically. B) Density of axis components on circular ExChrs. TOP: For each of the two ExChr-bearing strains depicted, the large images on the right show portions of nuclei from partially spread gonads prepared using conditions that preserve SCs, stained for HTP-3, COH-3/4 and SYP-1. At the left, schematic representations of the DNA composition of *meEx001* and *oxEx1578* are presented. BOTTOM: Quantification of axis foci on ExChrs in fully spread nuclei stained for either REC-8 or COH-3/4 together with HTP-3. In the graphs, each data point represents a single ExChr, with the X-axis indicating the numbers of REC-8 or COH-3/4 foci and the Y-axis indicating the numbers of HTP-3 foci counted on that ExChr. All data acquired are presented. For both ExChrs examined, the numbers of HTP-3 and REC-8 or COH-3/4 foci are strongly correlated (p<0.001) and the slopes of the linear regression lines are near to 1 (for *meEx001* REC-8 vs. HTP-3, r = .85, slope = .79; for *meEx001* COH-3/4 vs. HTP-3, r = .88, slope = .84; for *oxEx1578* REC-8 vs. HTP-3, r = .94, slope = .76; for *oxEx1578* COH-3/4 vs. HTP-3, r = .76, slope = .79). Further, in three of the four cases, a line with slope = 1 falls within the 95% confidence interval for the slope of the linear regression line (indicated by the gray shaded area), consistent with a near 1:1:1 correspondence between HORMAD, REC-8 and COH-3/4 foci, as observed for spread chromosome segments (depicted in FIG3). Thus, we infer that axis organization on the ExChrs is similar to that observed for normal chromosomes.

**Figure 2S1.**
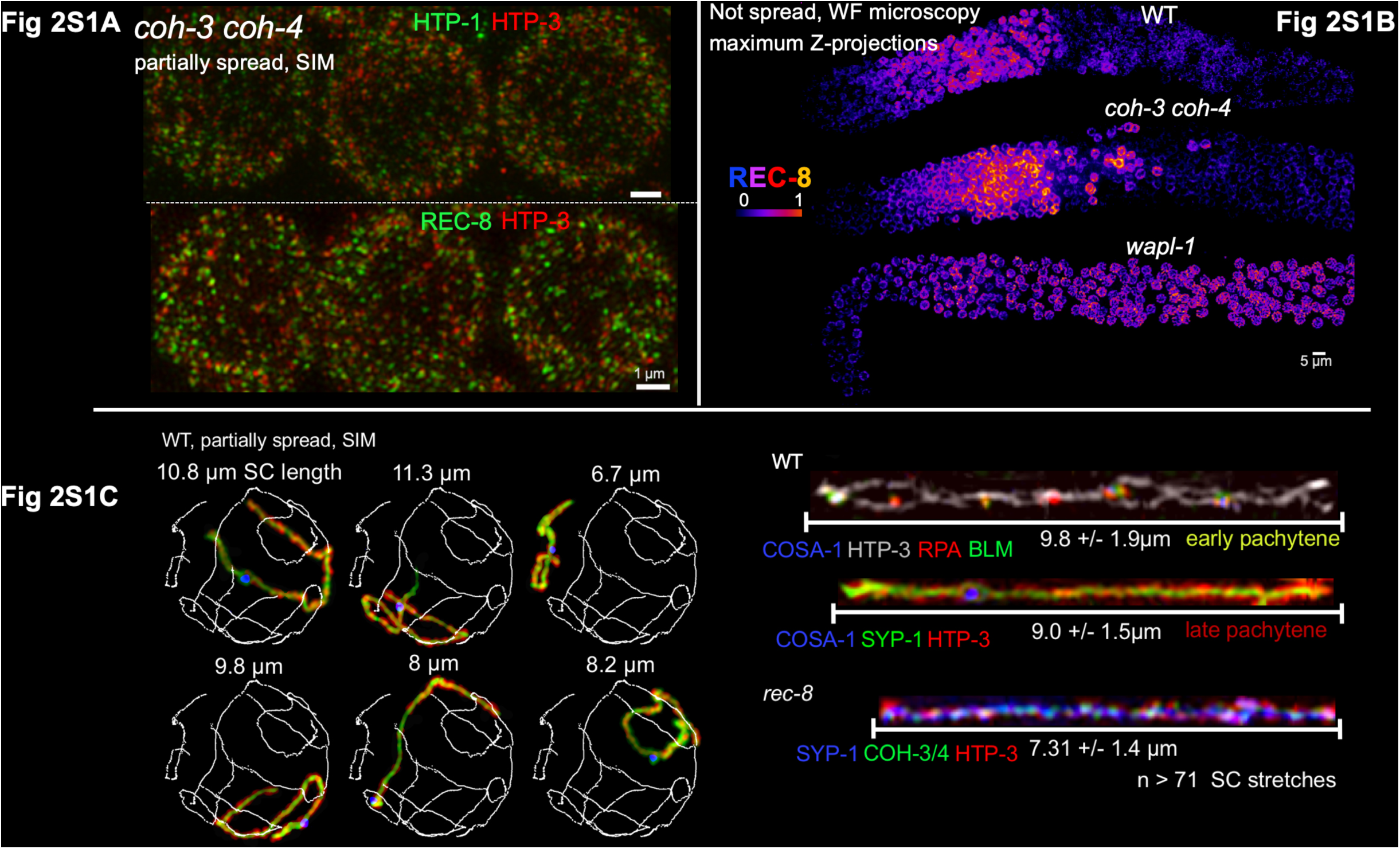
Functional distinctions between REC-8 and COH-3/4 cohesin complexes. A) SIM images of HORMAD proteins and REC-8 in partially spread meiotic prophase nuclei from the *coh-3 coh-4* mutant, illustrating the requirement for COH-3/4 in axis organization. HORMADs and REC-8 still bind to chromatin in this mutant, but the high degree of colocalization between HTP-3, HTP-1/2 and REC-8 seen in the WT is not observed and continuous axes do not form. B) Reduction in chromosome-associated REC-8 cohesion-conferring complexes occurs independently of installation of COH-3/4 axis-organizing cohesins. Non-spread gonads from WT, *coh-3/4* and *wapl-1* worms, with relative intensities of REC-8 immuno-fluorescence signals illustrated using the indicated color scale. The drop of REC-8 levels observed in the WT does not depend on the loading of COH-3/4, as it still occurs in their absence. However, a drop in REC-8 levels does not occur in the *wapl-1* mutant. C) SIM images of individual chromosomes from partially-spread meiotic prophase nuclei prepared using conditions that preserve association of SC central region proteins and maintain inter-axis distances comparable to *in situ* preparations (Woglar and Villeneuve, 2018). Left, Representative example images of SCs in a WT pachytene nucleus, provided together with traces of the paths of individual SCs that were generated using the ImageJ plugin “Simple Neurite Tracer” (Longair et al., 2011), illustrating that the paths of all six individual synapsed chromosome pairs can be reliably traced in 3D using this approach. Right, Example images of SCs of straightened chromosomes, indicating the average lengths of wild-type early pachytene SCs (identified based on the presence of multiple recombination foci [marked by RPA and BLM] per SC), wild-type late pachytene SCs (which have a single bright COSA-1-marked crossover site focus per SC), and the inter-sister SCs that are formed in a *rec-8* mutant.

**Figure 3S1.**
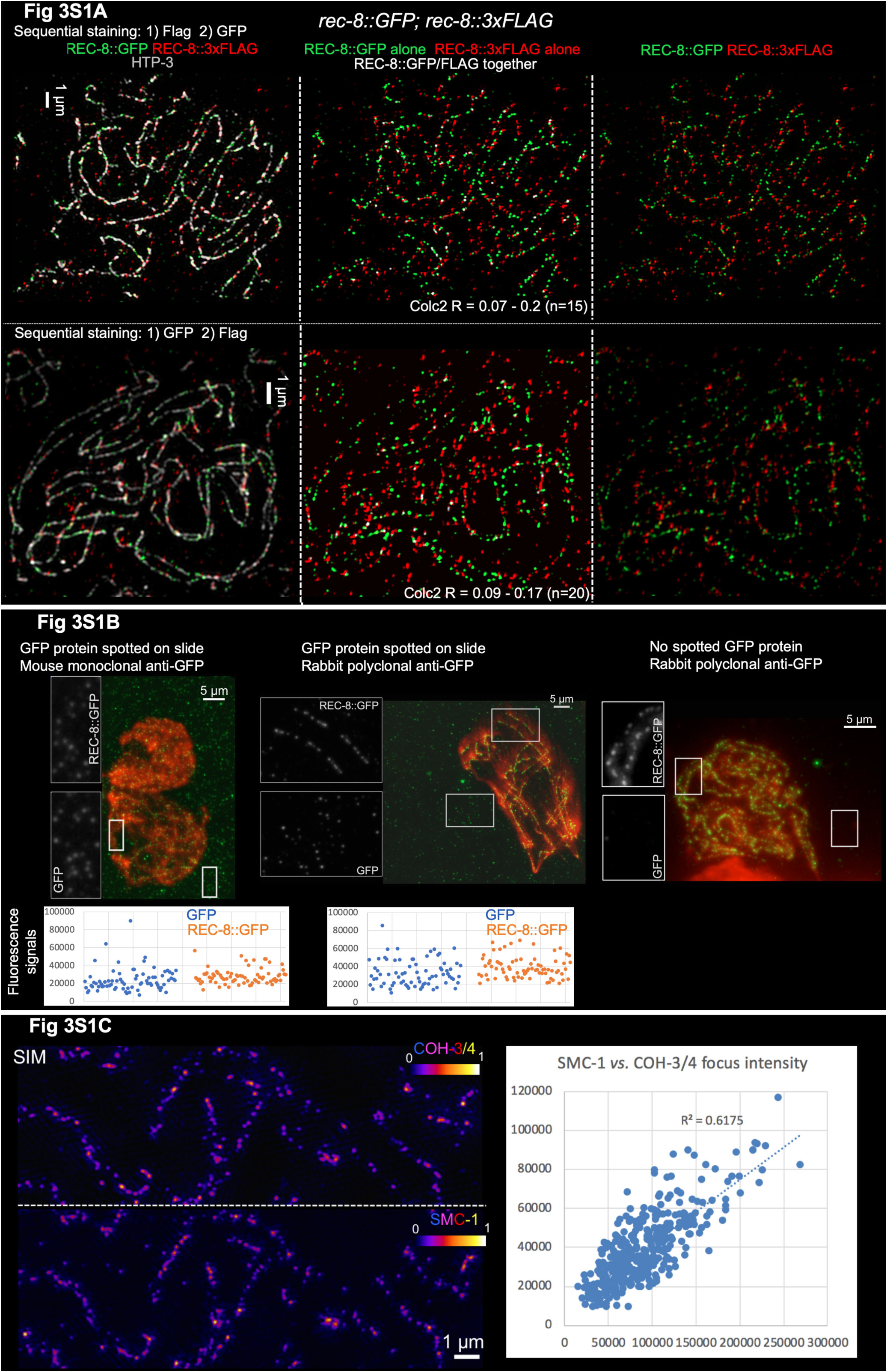
Supporting evidence for quantification of cohesin molecules. A) SIM images of fully-spread nuclei from worms expressing both REC-8::GFP and REC-8::3xFLAG, immuno-stained for HTP-3, FLAG and GFP using sequential immunostaining protocols. TOP: Anti-FLAG was applied first, followed by secondary detection of FLAG, before primary and secondary immuno-detection of HTP-3 and GFP. BOTTOM: Anti-GFP was applied first, followed by secondary detection of GFP, before primary and secondary immuno-detection of HTP-3 and FLAG. In all three orders in which the experiment was performed (sequential staining in both directions and simultaneous co-staining with all primary antibodies as in Figure 3A), GFP and FLAG signals both localize to chromosome axes (marked by HTP-3) in similar numbers, but most GFP and FLAG signals do not colocalize. Middle panels were generated using ImageJ plugin “ColocThreshold”, with white indicating the infrequent pixels with significant FLAG and GFP colocalization (colocalizing pixel are in white, non-colocalizing pixel are in red and green); contrast and brightness were adjusted for display to improve visibility of the white pixels. B) Similar immunofluorescence signals for purified GFP protein and chromosome-associated REC-8::GFP foci. Purified GFP protein was spotted onto a glass slide, and nuclei from worms expressing REC-8::GFP (as sole source of REC-8) were spread and fixed on top. Images are SUM-intensity projections of immunofluorescence detection of GFP with a mouse monoclonal antibody (left) or with a rabbit polyclonal antibody (center, right); images at the right are from a control slide without spotted GFP protein. Corresponding graphs plot background-subtracted fluorescence values for 4×4 pixel ROIs encompassing single REC-8::GFP foci (orange) or purified GFP protein foci (blue) from the same image. C) COH-3/4 and SMC-1 fluorescence values correlate with each other in fully spread nuclei from worms expressing SMC-1::HA. LEFT: SIM image of well-spread chromosome segments in which relative intensities of immuno-fluorescence signals are depicted using the indicated color scale, illustrating the strong similarity between the fluorescence intensity patterns for COH-3/4 (top) and SMC-1::HA (bottom). RIGHT: Graph plotting fluorescence values for SMC-1::HA (x-axis) and COH-3/4 (Y-axis) in 800×800 nm ROIs centered on individual axis foci from SUM projected, wide-field images. R^2^ value reflects a strong positive correlation between SMC-1 and COH-3/4 fluorescence values in individual foci.

**Figure 3S2.**
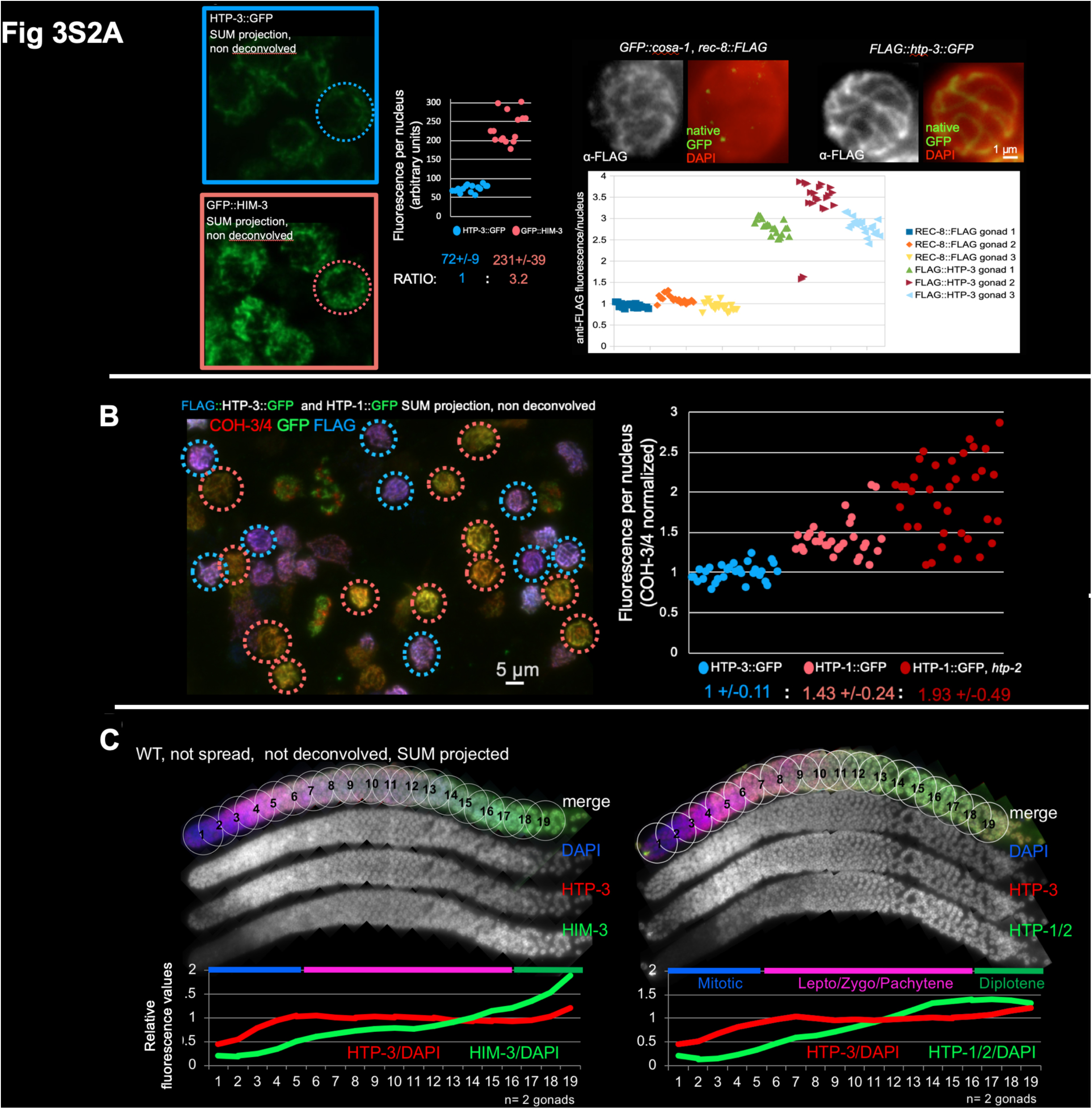
Assessment of relative levels of chromosome associated HORMAD proteins in whole nuclei. A) LEFT: SUM-projected non-deconvolved wide field images (left) of leptotene nuclei from partially-spread gonads from worms expressing either 3xFLAG::HTP-3::GFP or GFP::HIM-3, processed in parallel for anti-GFP immuno-fluorescence on the same slide. Each data point in the graph (right) represents the total fluorescence in a single nucleus. RIGHT: : SUM-projected non-deconvolved wide field images of late pachytene nuclei from partially-spread gonads from worms expressing either REC-8::3xFLAG and GFP::COSA-1 or 3xFLAG::HTP-3::GFP, processed in parallel for anti-FLAG immuno-fluorescence on the same slide; native GFP fluorescence was used to distinguish the genotypes of the gonads analyzed. Each data point in the graph represents the total fluorescence in a single nucleus; data are normalized to: Average REC-8::FLAG fluorescence = 1. B) Fully spread nuclei from worms expressing 3xFLAG::HTP-3::GFP and from worms expressing HTP-1::GFP (either in the presence or absence of HTP-2) were prepared as in FIG3D on the same slide (HTP-2 is present in the example image on the left). Nuclei were stained sequentially for GFP and COH-3/4, followed by detection of the FLAG epitope to identify the nuclei expressing 3xFLAG::HTP-3::GFP. Total immuno-fluorescence signals for GFP (green channel) and COH-3/4 (far-red channel) were measured in ROIs drawn around each individual nucleus. Each data point in the graph represents the ratio between total GFP signal and total COH-3/4 signal for a single nucleus (to account for differences in the degree of spreading of individual nuclei). Data for the two experiments were normalized to each other using the mean value for HTP-3::GFP as a normalization standard; In the graph, HTP-3::GFP values from the first experimental setup (presence of HTP-2 in both genotypes) are represented. C) SUM intensity projections of non-spread WT gonads stained for DAPI, HTP-3 and HTP-1/2, or DAPI, HTP-3 and HIM-3, and the gonads were divided as depicted into nineteen equal-sized, half-overlapping ROIs from the mitotic tip to start of cellularization (late diplotene). For each HORMAD being evaluated (HTP-3, HIM-3 or HTP-1/2), the ratio between total immuno-fluorescence signal and total DAPI signal was determined for each ROI. For the plots of relative fluorescence (for each HORMAD) over the course of meiotic prophase progression, measured values were normalized to the average value for the set of ROIs spanning from the onset of meiotic prophase (ROI 6) through the end of the pachytene stage (ROI 16-17). Two gonads were averaged for each staining. Note that HTP-3 levels remain stable over the course of meiotic prophase, whereas HTP-1/2 and HIM-3 levels increase.

**Figure 4S1.**
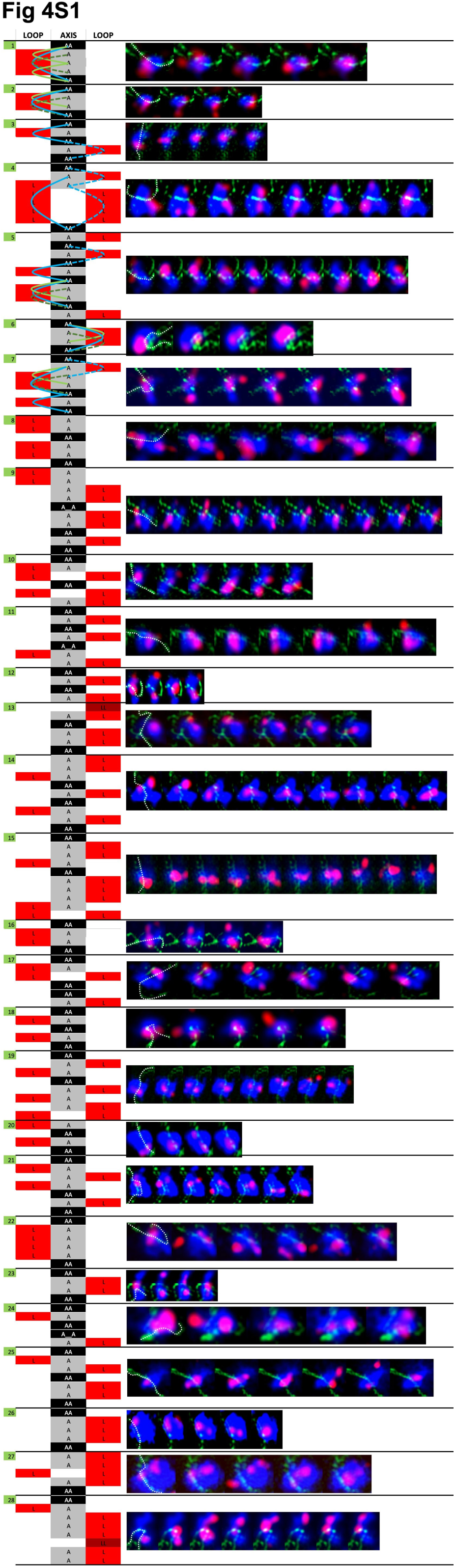
Display of individual hybridization sequences allowing inferences of DNA Loop paths. Each panel depicts a sequence of images (from left to right) from iterative FISH experiments in which consecutive 100 kbp DNA segments were individually visualized in red (using pools of secondary readout oligos) and the full 1 Mbp probed region was visualized in blue (fiducial probe). These panels depict a subset of chromosome segments of the analyzed data set that met the following criteria: 1) the two sister chromatid signals could be resolved for at least every other step, and 2) the fiducial probe signal could be assigned unambiguously to a chromosome axis, visualized by HTP-3 immunofluorescence. For reference, a dashed white line in the initial frame of each panel indicates the path of the HTP-3 axis signal considered for analysis. When axes were imaged from a frontal perspective, *i.e*. when two roughly parallel axis signals could be detected, the fiducial probe signal was inferred to correspond to DNA associated with the closer of the two axis signals. Ambiguous signals (which can represent the majority in a given microscopic field) were not analyzed. Panels 1 and 2 correspond to two different portions of the same chromosome. For a few individual hybridization steps in sequences 7, 8, 24 and 27, complex fluorescence signals were co-assigned to a given chromatid based on proximity. Sequence 4 includes one ambiguous step (step two) that can be interpreted differently than displayed. On the left, the consecutive 100 kbp DNA segments represented in each left-to-right image panel are schematically represented from top to bottom. Positions where FISH signals overlap with the axis signal (A) are indicated in the center column in gray or black cells, while positions where FISH signals are in loops >100 kbp in size (L) are indicated in red cells flanking the center column. Unresolvable signals that colocalized with the assigned chromosome axis are represented as “AA” (63 cases in the displayed data set). If signals were resolvable, but both colocalize with the assigned chromosome axis, they are represented as “A A” (3 cases). If signals were not resolvable but did not colocalize with the chromosome axis, they are represented as LL (2 cases, dark red fields). For the first seven chromosome segments analyzed, curves illustrating potential paths of DNA loops that are larger than 100 kbp (*i.e*. where one or several consecutive 100 kbp DNA segments were scored as “L”) are overlaid on the schematic diagrams. In this representation, loops from two different sister chromatids are indicated by solid and dashed lines, respectively. In some cases, *e.g*. panel 3, only one set of potential paths is compatible with the underlying diagram and corresponding images. For other cases, alternative paths are possible; the maximum-sized potential loop paths are indicated in blue, minimum-sized potential loop paths in green. We also analyzed the full 10-step FISH sequences for a total of 26 chromosomes, including 13 of the 27 chromosomes represented here plus an additional 13 chromosomes. This unbiased data set was used for Figure 4C-D and the corresponding statistical analyses presented in the text. Notably, LL signals represent only a small fraction (3.5%) of the total signals in this data set; thus, the conclusion that symmetric loops >100 kbp in size are infrequent would not be affected by the possibility that unresolved LL signals might be underrepresented among the displayed images.

**Figure 4S2.**
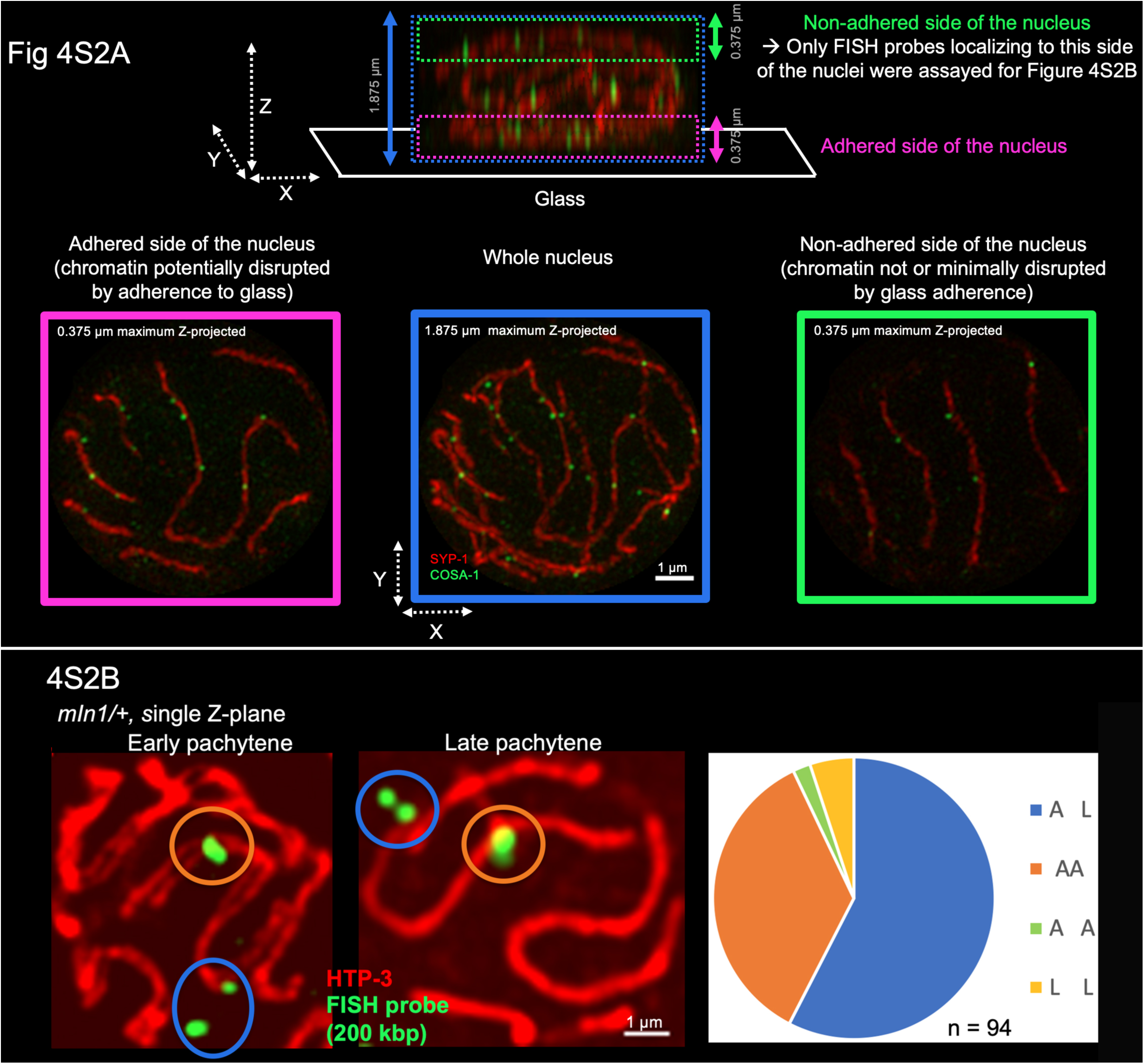
Testing asymmetry of sister chromatid lops in minimally disrupted tissue. A) Illustration indicating which side of the nucleus was assayed to determine the distances between FISH signal junctions and the chromosome axis for the analysis in FIG4S2B. The side of the nucleus that adheres to the glass (magenta frame, left) in the partial spreading procedure is visibly more flattened than the non-adhering side (green frame, right). Moreover, adherence of chromatin/DNA to the glass slide itself may potentially introduce artifacts affecting DNA/chromatin organization. Therefore, only FISH signals that were associated with chromosome segments in the non-adhering side of the nucleus were assayed for FIG4S2B. B) Immuno-FISH evidence for asymmetry between sister chromatid loops in minimally disrupted tissue in 3D. *mIn1*, a rearranged version of chromosome II harboring an 8.2 Mbp internal inversion was used for assessment of sister chromatid loop relationships. *mIn1* heterozygosity results in wide separation between the FISH signals for the normal and rearranged homologs, thereby enabling assessment of sister chromatid signals. Images show two examples (early pachytene, left and late pachytene, right) of single Z-planes in which both 200 kbp FISH signals (chromosome II.. 11.5-11.7 Mbp) could be found in the same Z-plane. FISH signals were only assessed if they were visible as a single focus or two foci; splintered signals were excluded. The distance between the centroid of each FISH signal and the center of the nearest axis signal (visualized by HTP-3 immunostaining) was measured in 3D. A signal was classified as “A” if its centroid was located < 240 nm (< 3 pixel in XY) from the center of the nearest axis segment and classified as “L” if located farther away.

